# Breadth and function of antibody response to acute SARS-CoV-2 infection in humans

**DOI:** 10.1101/2020.08.28.267526

**Authors:** Kuan-Ying A. Huang, Tiong Kit Tan, Ting-Hua Chen, Chung-Guei Huang, Ruth Harvey, Saira Hussain, Cheng-Pin Chen, Adam Harding, Javier Gilbert-Jaramillo, Xu Liu, Michael Knight, Lisa Schimanski, Shin-Ru Shih, Yi-Chun Lin, Chien-Yu Cheng, Shu-Hsing Cheng, Yhu-Chering Huang, Tzou-Yien Lin, Jia-Tsrong Jan, Che Ma, William James, Rodney S. Daniels, John W. McCauley, Pramila Rijal, Alain R. Townsend

## Abstract

Serological and plasmablast responses and plasmablast-derived IgG monoclonal antibodies (MAbs) have been analysed in three COVID-19 patients with different clinical severities. Potent humoral responses were detected within 3 weeks of onset of illness in all patients and the serological titre was elicited soon after or concomitantly with peripheral plasmablast response. An average of 13.7% and 13.0% of plasmablast-derived MAbs were reactive with virus spike glycoprotein or nucleocapsid, respectively. A subset of anti-spike (10 of 32) and over half of anti-nucleocapsid (19 of 35) antibodies cross-reacted with other betacoronaviruses tested and harboured extensive somatic mutations, indicative of an expansion of memory B cells upon SARS-CoV-2 infection. Fourteen of 32 anti-spike MAbs, including five anti-RBD, three anti-non-RBD S1 and six anti-S2, neutralised wild-type SARS-CoV-2 in independent assays. Anti-RBD MAbs were further grouped into four cross-inhibiting clusters, of which six antibodies from three separate clusters blocked the binding of RBD to ACE2 and five were neutralising. All ACE2-blocking anti-RBD antibodies were isolated from two patients with prolonged fever, which is compatible with substantial ACE2-blocking response in their sera. At last, the identification of non-competing pairs of neutralising antibodies would offer potential templates for the development of prophylactic and therapeutic agents against SARS-CoV-2.

## Introduction

In late 2019, a novel coronavirus emerged and was identified as the cause of a cluster of respiratory infection cases in Wuhan, China. It spread quickly around the world. In March of 2020 a pandemic was declared by the World Health Organization, the virus was formally named as Severe Acute Respiratory Syndrome Coronavirus 2 (SARS-CoV-2) and the resulting disease was named COVID-19. As of 1 October 2020, there have been over 33 million confirmed cases of SARS-CoV-2 infection with 1,010,986 deaths (World Health Organization, https://covid19.who.int/).

There is no fully effective drug or licenced vaccine for COVID-19. Antibodies neutralise SARS-CoV-2 *in vitro,* offering hope that a vaccine may induce a protective response, and antibodies may be an effective treatment for COVID-19 in clinical practice. Convalescent plasma is being tested in ongoing clinical trials as a therapy for COVID-19 (1, 2), and was previously used in the treatment of SARS (3). The virus spike glycoprotein is a target of neutralising antibodies, which makes it a key candidate for both vaccine development and immunotherapy (4). B cell responses in COVID-19 patients have been detected concomitantly with follicular helper T cell responses from week one after illness onset (5). In SARS patients, B cell responses typically arise first against the nucleocapsid protein then, within four to eight days after symptom onset, antibody responses to spike glycoprotein have been found; neutralising antibody responses begin to develop by week two, and most patients develop neutralising antibodies by week three (6). Two serological studies of COVID-19 patients showed anti-SARS-CoV-2 IgG seroconversion at week three after onset and some cross-reactivity to nucleocapsid of SARS (7, 8).

Antibodies may play a role in protection against SARS-CoV-2 infection. The underlying B-cell response leading to the rapid production of plasmablasts (antibody-secreting cells) that secrete antibodies upon natural SARS-CoV-2 exposure/infection is only beginning to be understood (5). Here, we characterised the infection-induced serological and plasmablast responses and the derived IgG anti-SARS-CoV-2 spike glycoprotein and nucleocapsid monoclonal antibodies (MAbs) from adult patients with laboratory-confirmed COVID-19. The antigenic specificity and breadth of antibodies and the sequence of their variable domains have been characterised in detail. Virus neutralising antibodies were detected that bound epitopes on receptor-binding domain (RBD), non-RBD regions of the S1 polypeptide, and the S2 polypeptide of the spike glycoprotein.

## Results

### Serological response and anti-spike glycoprotein and anti-nucleocapsid IgG antibodies from circulating plasmablasts

Serum IgG antibodies to the virus spike glycoprotein and the isolated RBD were analysed by indirect enzyme-linked immunosorbent and flow cytometry assays in three patients with laboratory-confirmed COVID-19. The clinical characteristics of the three patients studied are shown in Supplemental Table 1. Antibodies to spike glycoprotein and RBD were detected in all three patients after week 3 of illness onset (Figure 1a). Case A showed a robust response to spike glycoprotein and the isolated RBD by day (D) 22 (D22). Longitudinal sera from case B and C showed lower anti-spike glycoprotein and RBD IgG titres at week 1 or the beginning of week 2 and an elevated titre that peaked at the end of week 2 through week 3 (a peak 50% effective dilution (ED_50_) titre to RBD 1:1,051 at D18 in case B and 1:588 at D14 in case C). Case B had prolonged fever and developed pneumonia at the end of week 2, which was followed by a robust increase of anti-spike glycoprotein and RBD IgG titres at week 3 (Figure 1a, Supplemental Table 1). By contrast, case C experienced a two-day course of febrile illness and reduction of all symptoms within the first week, followed by anti-spike glycoprotein and RBD IgG titres that peaked earlier at the end of week 2 (Figure 1a, Supplemental Table 1).

**Figure 1.**
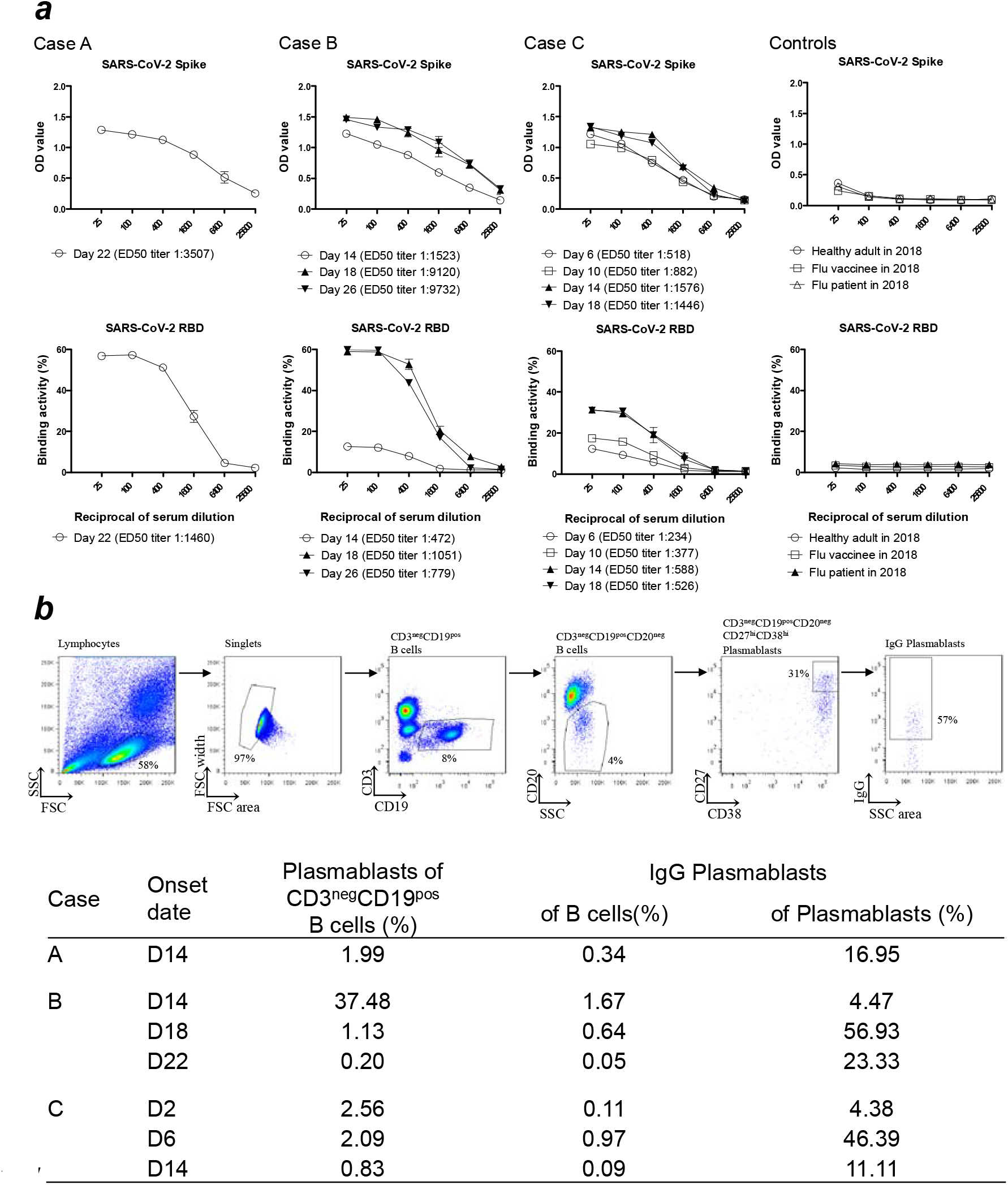
The IgG serology and plasmablast response to acute SARS-CoV-2 infection among enrolled patients. (a) The binding activity of post-infection sera IgG with SARS-CoV-2 spike glycoprotein in an ELISA and SARS-CoV-2 RBD assessed by flow cytometry on transfected cells, among enrolled patients. Each experiment was repeated twice. Values are presented as mean ± standard error of the mean. Two sera from healthy adults (one collected at day 28 post 2018-19 influenza vaccination and one collected from an influenza-infected patient 9 days after symptom onset) in 2018 were included as controls. Linear regression was used to determine the 50% end-point dilution (ED_50_). (b) The gating strategy used for peripheral total B cells, plasmablasts and IgG plasmablasts in flow cytometry. The frequency of circulating plasmablasts (percentage of total B cells) among enrolled cases was measured by flow cytometry. Onset date (D = Day).

An increased frequency of circulating plasmablasts was detected in all three patients (healthy adults baseline less than 1%) (5) (Figure 1b). In case A, a plasmablast response containing a substantial IgG subset was detected at the end of week 2 (D14), followed by a high serological titre at week 3 (D22). Case B had a robust plasmablast response at the end of week 2 (D14) but the IgG plasmablast subset continued to rise, dominated at week 3 (D18), but then subsided at the beginning of week 4 (D22), which is compatible with high anti-spike glycoprotein and RBD IgG serological titres at week 3. Case C produced a significant early plasmablast response at the end of week 1 (D6), the IgG plasmablast subset dominated at the same time, and both the plasmablast response and its IgG subsets subsided at the end of week 2 (D14).

Circulating plasmablasts were identified and used to generate human IgG monoclonal antibodies (MAbs) from the three patients (Figure 2a). A total of 219 plasmablast-derived IgG MAbs were produced, of which 67 (10 of 50 from case A, 48 of 131 from case B, 9 of 38 from case C) were shown to bind spike glycoprotein or nucleocapsid antigens of SARS-CoV-2 by one or more of the following: staining of spike glycoprotein and RBD-expressing cells, ELISA, or immunofluorescence and specific virus neutralisation. Averages of 13.7±6.8% (6.0-18.4%) and 13.0±7.3% (5.3-19.9%) of plasmablast-derived IgG MAbs were reactive with virus spike glycoprotein and nucleocapsid, respectively.

**Figure 2.**
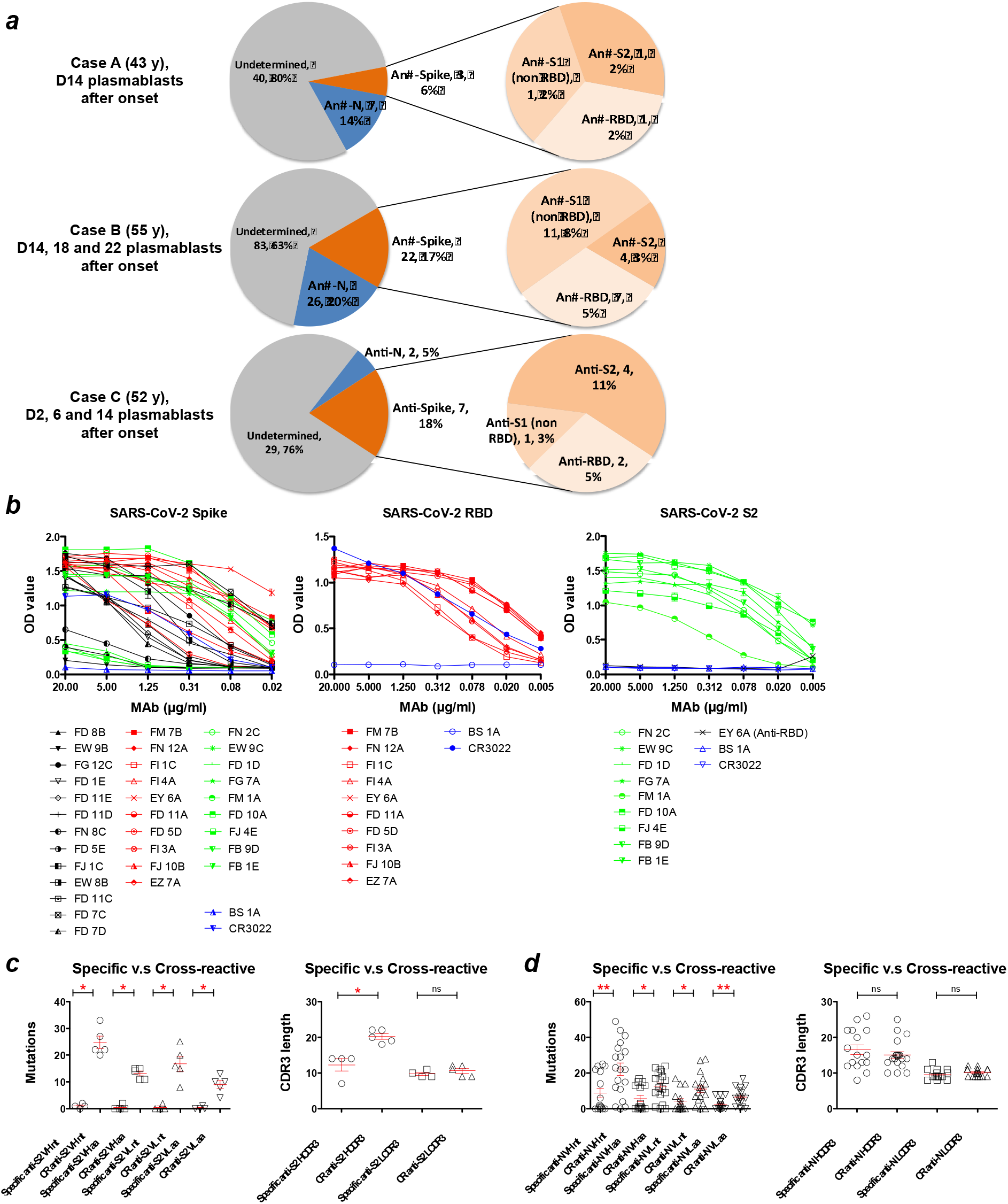
Plasmablast-derived IgG monoclonal antibodies from three COVID-19 patients. (a) A total of 219 IgG monoclonal antibodies were produced from COVID-19 patients (50 from case A, 131 from case B, 38 from case C). An average of 13.7±6.8% and 13.0±7.3% of antibodies were reactive with spike glycoprotein (S) and nucleocapsid (N) antigens of SARS-CoV-2, respectively. The data are presented as specificity, number of antibodies, and the percentage of total antibodies isolated from each patient. (b) The binding activity of anti-SARS-CoV-2 MAbs with spike glycoprotein, RBD and the S2 subunit in ELISA. Anti-influenza H3 MAb BS-1A and anti-SARS RBD CR3022 were included as controls. Each experiment was repeated twice. The OD_450_ values are presented as mean ± standard error of the mean. Panels (c) and (d) show numbers of variable domain mutations in MAb genes and variation in MAb CDR3 lengths among anti-S2 and anti-N MAbs, respectively. Antibodies that strongly cross-react with at least one betacoronavirus (SARS or MERS or OC43) were defined as cross-reactive MAbs. CDR3 length and mutation numbers are presented as mean ± standard error of the mean (anti-S2, specific, n=4 versus cross-reactive, n=5; anti-N, specific, n=16 versus cross-reactive, n=19). The two-tailed Mann-Whitney test was performed to compare the mutations between two groups. * p < 0.05, ** p < 0.01 ; D, =Day ; ns, non-significant ; CR, cross-reactive.

### Genetic and phenotypic characteristics of anti-spike glycoprotein antibodies

Among 32 anti-spike MAbs, 10 bound to the RBD, 13 to non-RBD S1, and the other 9 to the S2 subunit of the SARS-CoV-2 spike glycoprotein (Figure 2a and Table 1). Twenty-four of these MAbs bound to virus antigens as assessed by immunofluorescence of SARS-CoV-2-infected Vero E6 cells (Supplemental Figure 1), suggesting that the majority of anti-spike glycoprotein human antibodies recognise complex conformational epitopes on the virus glycoprotein. Moderate binding was observed for a subset (4 of 9) of anti-S2 MAbs with full-length spike glycoprotein ectodomain in the indirect ELISA but bound strongly to the isolated S2 subunit (Figure 2b).

Ten of 32 anti-SARS-CoV-2 spike glycoprotein MAbs cross-reacted with the glycoproteins of other betacoronaviruses, including SARS, MERS or human common cold coronavirus OC43 in ELISA (Table 1), suggesting the presence of conserved epitopes on the spike glycoproteins of betacoronaviruses.

**Table 1.**
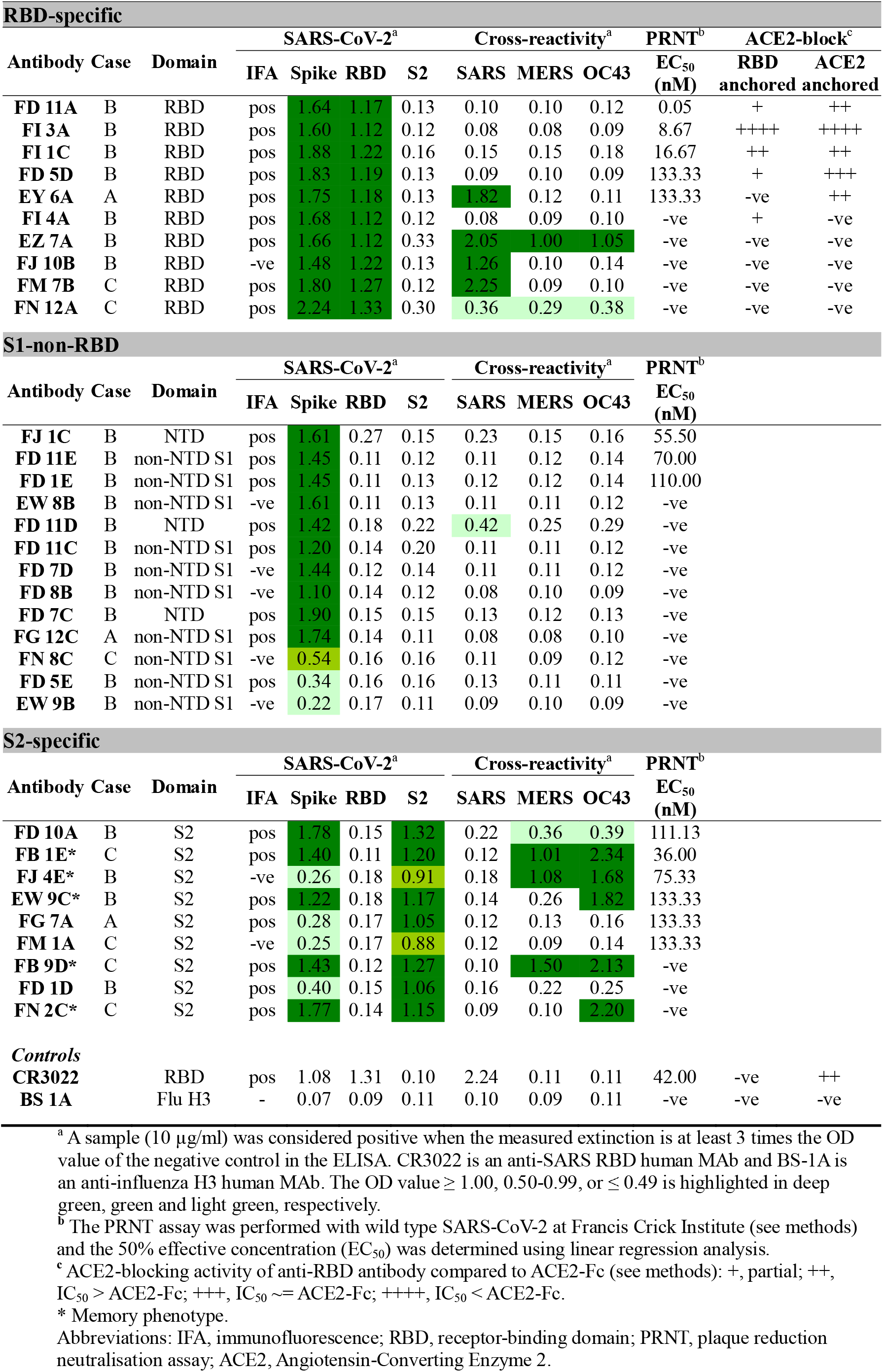
The antigenic specificity, cross-reactivity and function of 32 anti-SARS-CoV-2 spike antibodies derived from COVID-19 patients.

| RBD-specific |  |  |  |  |  |  |  |  |  |  |  |  |
| --- | --- | --- | --- | --- | --- | --- | --- | --- | --- | --- | --- | --- |
| Antibody | Case | Domain | SARS-CoV-2<sup>a</sup> |  |  |  | Cross-reactivity<sup>a</sup> |  |  | PRNT<sup>b</sup> | ACE2-block<sup>c</sup> |  |
|  |  |  | IFA | Spike | RBD | S2 | SARS | MERS | OC43 | EC<sub>50</sub><br>(nM) | RBD<br>anchored | ACE2<br>anchored |
| FD 11A | B | RBD | pos | 1.64 | 1.17 | 0.13 | 0.10 | 0.10 | 0.12 | 0.05 | + | ++ |
| FI 3A | B | RBD | pos | 1.60 | 1.12 | 0.12 | 0.08 | 0.08 | 0.09 | 8.67 | ++++ | ++++ |
| FI 1C | B | RBD | pos | 1.88 | 1.22 | 0.16 | 0.15 | 0.15 | 0.18 | 16.67 | ++ | ++ |
| FD 5D | B | RBD | pos | 1.83 | 1.19 | 0.13 | 0.09 | 0.10 | 0.09 | 133.33 | + | +++ |
| EY 6A | A | RBD | pos | 1.75 | 1.18 | 0.13 | 1.82 | 0.12 | 0.11 | 133.33 | -ve | ++ |
| FI 4A | B | RBD | pos | 1.68 | 1.12 | 0.12 | 0.08 | 0.09 | 0.10 | -ve | + | -ve |
| EZ 7A | B | RBD | pos | 1.66 | 1.12 | 0.33 | 2.05 | 1.00 | 1.05 | -ve | -ve | -ve |
| FJ 10B | B | RBD | -ve | 1.48 | 1.22 | 0.13 | 1.26 | 0.10 | 0.14 | -ve | -ve | -ve |
| FM 7B | C | RBD | pos | 1.80 | 1.27 | 0.12 | 2.25 | 0.09 | 0.10 | -ve | -ve | -ve |
| FN 12A | C | RBD | pos | 2.24 | 1.33 | 0.30 | 0.36 | 0.29 | 0.38 | -ve | -ve | -ve |

| S1-non-RBD |  |  |  |  |  |  |  |  |  |  |
| --- | --- | --- | --- | --- | --- | --- | --- | --- | --- | --- |
| Antibody | Case | Domain | SARS-CoV-2<sup>a</sup> |  |  |  | Cross-reactivity<sup>a</sup> |  |  | PRNT<sup>b</sup> |
|  |  |  | IFA | Spike | RBD | S2 | SARS | MERS | OC43 | EC<sub>50</sub><br>(nM) |
| FJ 1C | B | NTD | pos | 1.61 | 0.27 | 0.15 | 0.23 | 0.15 | 0.16 | 55.50 |
| FD 11E | B | non-NTD S1 | pos | 1.45 | 0.11 | 0.12 | 0.11 | 0.12 | 0.14 | 70.00 |
| FD 1E | B | non-NTD S1 | pos | 1.45 | 0.11 | 0.13 | 0.12 | 0.12 | 0.14 | 110.00 |
| EW 8B | B | non-NTD S1 | -ve | 1.61 | 0.11 | 0.13 | 0.11 | 0.11 | 0.12 | -ve |
| FD 11D | B | NTD | pos | 1.42 | 0.18 | 0.22 | 0.42 | 0.25 | 0.29 | -ve |
| FD 11C | B | non-NTD S1 | pos | 1.20 | 0.14 | 0.20 | 0.11 | 0.11 | 0.12 | -ve |
| FD 7D | B | non-NTD S1 | -ve | 1.44 | 0.12 | 0.14 | 0.11 | 0.11 | 0.12 | -ve |
| FD 8B | B | non-NTD S1 | -ve | 1.10 | 0.14 | 0.12 | 0.08 | 0.10 | 0.09 | -ve |
| FD 7C | B | NTD | pos | 1.90 | 0.15 | 0.15 | 0.13 | 0.12 | 0.13 | -ve |
| FG 12C | A | non-NTD S1 | pos | 1.74 | 0.14 | 0.11 | 0.08 | 0.08 | 0.10 | -ve |
| FN 8C | C | non-NTD S1 | -ve | 0.54 | 0.16 | 0.16 | 0.11 | 0.09 | 0.12 | -ve |
| FD 5E | B | non-NTD S1 | pos | 0.34 | 0.16 | 0.16 | 0.13 | 0.11 | 0.11 | -ve |
| EW 9B | B | non-NTD S1 | -ve | 0.22 | 0.17 | 0.11 | 0.09 | 0.10 | 0.09 | -ve |

| S2-specific |  |  |  |  |  |  |  |  |  |  |
| --- | --- | --- | --- | --- | --- | --- | --- | --- | --- | --- |
| Antibody | Case | Domain | SARS-CoV-2<sup>a</sup> |  |  |  | Cross-reactivity<sup>a</sup> |  |  | PRNT<sup>b</sup> |
|  |  |  | IFA | Spike | RBD | S2 | SARS | MERS | OC43 | EC<sub>50</sub><br>(nM) |
| FD 10A | B | S2 | pos | 1.78 | 0.15 | 1.32 | 0.22 | 0.36 | 0.39 | 111.13 |
| FB 1E* | C | S2 | pos | 1.40 | 0.11 | 1.20 | 0.12 | 1.01 | 2.34 | 36.00 |
| FJ 4E* | B | S2 | -ve | 0.26 | 0.18 | 0.91 | 0.18 | 1.08 | 1.68 | 75.33 |
| EW 9C* | B | S2 | pos | 1.22 | 0.18 | 1.17 | 0.14 | 0.26 | 1.82 | 133.33 |
| FG 7A | A | S2 | pos | 0.28 | 0.17 | 1.05 | 0.12 | 0.13 | 0.16 | 133.33 |
| FM 1A | C | S2 | -ve | 0.25 | 0.17 | 0.88 | 0.12 | 0.09 | 0.14 | 133.33 |
| FB 9D* | C | S2 | pos | 1.43 | 0.12 | 1.27 | 0.10 | 1.50 | 2.13 | -ve |
| FD 1D | B | S2 | pos | 0.40 | 0.15 | 1.06 | 0.16 | 0.22 | 0.25 | -ve |
| FN 2C* | C | S2 | pos | 1.77 | 0.14 | 1.15 | 0.09 | 0.10 | 2.20 | -ve |
| Controls |  |  |  |  |  |  |  |  |  |  |  |  |
| CR3022 |  | RBD | pos | 1.08 | 1.31 | 0.10 | 2.24 | 0.11 | 0.11 | 42.00 | -ve | ++ |
| BS 1A |  | Flu H3 | - | 0.07 | 0.09 | 0.11 | 0.10 | 0.09 | 0.11 | -ve | -ve | -ve |

<sup>a</sup> A sample (10 µg/ml) was considered positive when the measured extinction is at least 3 times the OD value of the negative control in the ELISA. CR3022 is an anti-SARS RBD human MAb and BS-1A is an anti-influenza H3 human MAb. The OD value $\geq 1.00$ , 0.50-0.99, or $\leq 0.49$ is highlighted in deep green, green and light green, respectively.
<sup>b</sup> The PRNT assay was performed with wild type SARS-CoV-2 at Francis Crick Institute (see methods)
and the 50% effective concentration ( $EC_{50}$ ) was determined using linear regression analysis. <sup>c</sup> ACE2-blocking activity of anti-RBD antibody compared to ACE2-Fc (see methods): +, partial; ++, $IC_{50} > ACE2-Fc$ ; +++, $IC_{50} \sim ACE2-Fc$ ; +++++, $IC_{50} < ACE2-Fc$ . \* Memory phenotype. Abbreviations: IFA, immunofluorescence; RBD, receptor-binding domain; PRNT, plaque reduction neutralisation assay; ACE2, Angiotensin-Converting Enzyme 2.

Each of 32 anti-spike glycoprotein MAbs was encoded by a unique set of heavy chain VDJ and light chain VJ rearrangements in the variable domain (Supplemental Table 2). Fourteen of 32 SARS-CoV-2 spike-reactive MAb genes possessed low numbers of somatic mutations resulting in 0 or 1 amino acid substitutions suggesting a *de novo* B cell response to the SARS-CoV-2 virus in humans. Six of the 32 MAb genes possessed ≥ 20 nucleotide mutations, and these cross-reacted on other beta-coronaviruses, including OC43. Of the nine anti-S2 antibodies five cross-reacted on OC43 virus and three of these also cross-reacted on MERS (Table 1). All five cross-reactive anti-S2 antibodies had high rates of somatic mutation (25±5), indicating a memory phenotype, and three of the five were neutralising to a moderate level (half maximal effective concentration, EC_50_, 36-133.33 nM, Table 1).

The CDR3 length varied among anti-spike glycoprotein antibodies (Supplemental Table 2). No significant differences were found between anti-S2 and anti-S1 or anti-RBD subsets. Among anti-S2 MAbs, a significantly longer heavy chain CDR3 length was found in the cross-reactive group compared to the specific group (Cross-reactive 20±2 versus Specific 12±4, p= 0.02, two-tailed Mann-Whitney test; Figure 2c), indicating that a long CDR3 may play a role in antigen binding, which is also found in several broadly reactive human MAbs against human immunodeficiency virus and influenza virus (9, 10).

The binding activities of 10 anti-RBD MAbs were further characterised in detail. Using MDCK-SIAT1 cells transduced to express the RBD and flow cytometry, binding activities of the anti-RBD MAbs were shown to vary with 50% binding concentration from 0.10 to 1.83 μg/ml (Supplemental Figure 2). The MAbs with strong anti-RBD binding have a relatively long heavy chain CDR3 length (50% binding concentration <0.5 μg/ml versus >0.5 μg/ml, p=0.03, two-tailed Mann-Whitney test; Supplemental Figure 3).

### Neutralisation by anti-spike glycoprotein antibodies

The optimum method for detecting neutralisation of SARS-CoV-2 has not been established. For example, different laboratories have obtained varying results for the well-defined monoclonal antibody CR3022 (11–13). We have therefore obtained results from neutralisation assays performed in independent laboratories : a systematic survey of the thirty two anti-spike MAbs at the Francis Crick Institute (London), with follow up for a selection of MAbs at the Sir William Dunn School (Oxford) and Chang Gung Memorial Hospital (Taiwan).

The 32 anti-spike glycoprotein MAbs were systematically examined by plaque reduction neutralisation (PRNT) assay for neutralisation of wild type SARS-CoV-2 virus (see methods; summarised in Table 1). A total of 14 neutralising antibodies distributed between different regions of the spike glycoprotein were identified: 5 of 10 to RBD, 3 of 13 to S1 (non-RBD), 6 of 9 to S2. The EC50 concentrations, as a measure of potency, ranged from 0.05 to ~133 nM (8 ng/ml - ~20 μg/ml).

Neutralisation of anti-RBD antibodies was corroborated by a microneutralisation test that measured a reduction in fluorescent focus-forming units (see methods, Supplemental Figure 4). MAbs were also tested by a PCR-based neutralisation assay (see methods): inhibition of virus replication was measured by quantitative PCR in the supernatant bathing the infected cells. This results corroborated that anti-RBD FD 11 A, anti-RBD FI 3A, anti-RBD FD 5D, anti-RBD EY 6A and anti-S2 EW 9C, as crude culture supernatants, reduced the virus signal from ~56- to ~10,085-fold (Supplemental Figure 5).

### ACE2 blockade by anti-RBD antibodies

Potent neutralising antibodies to the RBD of SARS-CoV-2 spike glycoprotein were identified and we thus analyse the blockade of the ACE2-RBD interaction by anti-RBD antibodies in two assays (Figure 3, Table 1), including an assay that we reported previously (14, 15). In the first assay unlabelled MAbs in at least 10-fold excess were mixed with biotin labelled ACE2-Fc and binding to the RBD, displayed on RBD-VLP (16) bound to an ELISA plate (Figure 3a, Supplemental Figure 6). Anti-RBD neutralising antibodies FI 3A and FI 1C, as well as the unlabelled ACE2-Fc, strongly inhibited the binding of labelled ACE2-Fc to RBD. A partial inhibition was detected by anti-RBD antibodies FD 11A, FD 5D and FI 4A. Another unlabelled nanobody VHH72-Fc in excess inhibited the binding of labelled ACE2-Fc to RBD in this assay. The structure of VHH72-Fc bound to RBD is known (17) and its footprint on the RBD does not overlap that of ACE2, so inhibition is thought to occur by steric hindrance.

**Figure 3.**
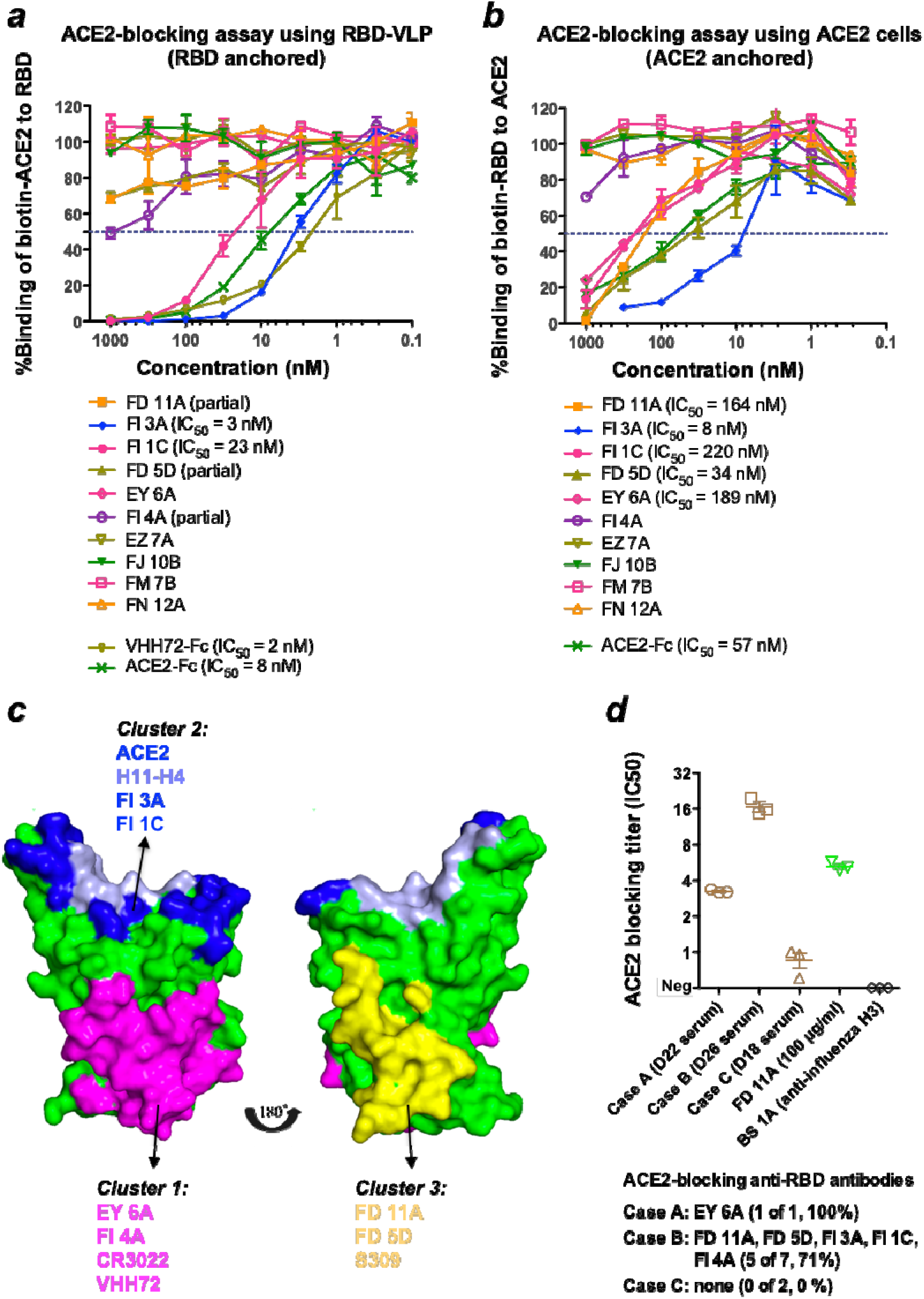
ACE2-blocking activities with anti-RBD antibodies and convalescent sera. The analyses were performed with **(a)** RBD anchored and **(b)** ACE2 anchored on plates (see methods). Anti-SARS-CoV-2 RBD nanobody VHH72 linked to the hinge and Fc region of human IgG1 and ACE2-Fc were included as controls. Experiments were performed in duplicate and repeated twice. **(c)** Mapping of neutralising anti-RBD antibodies on the SARS-CoV-2 RBD structure (PDB 6ZCZ) based on competitive binding and ACE2-blocking analyses. The RBD was colored in green. The epitopes recognized by EY 6A, CR3022 and VHH72 (cluster 1 MAb) (11, 15, 17) were colored in magenta. The epitopes recognized by ACE2 and H11-H4 (cluster 2 MAb) (14) were overlapping and colored in blue and light blue. The epitopes recognized by S309 (cluster 3 MAb) (19) were colored in yellow. **(d)** Convalescent sera were analysed in the ACE2-blocking (ACE2 anchored) assay. Experiments were performed in triplicate. Anti-RBD antibody FD 11A and anti-influenza H3 antibody BS 1A were included as controls. Data are presented as mean ± standard error of the mean. IC_50_, 50% inhibitory concentration.

In the second assay, we employed MDCK-SIAT1 cells overexpressing full-length human ACE2 as a transmembrane protein. Unlabelled antibodies or ACE2-Fc were mixed in excess with biotinylated RBD, and binding of RBD was detected with Streptavidin-HRP in ELISA (Figure 3b). The results of this assay mostly mirrored those of the first assay and confirmed that in this orientation anti-RBD neutralising antibodies FD 11A and FD 5D competed in excess with soluble RBD for binding to ACE2 (Figure 3b). In addition, anti-RBD neutralising antibody EY 6A competed with RBD for ACE2 binding. The binding pattern of EY 6A is analogous to a previously described antibody CR3022 (Table 1) (11). These two antibodies are known to bind to the same region of RBD away from the ACE2 binding site, but they influence the binding kinetics of RBD to ACE2, presumably through steric effects (15).

### Division of anti-RBD antibodies into cross-inhibiting groups

The ten anti-RBD MAbs were then divided into cross-inhibiting groups as described for human MAbs to Ebola (18) by assessing competition of unlabelled antibodies at 10-fold (or greater) excess over a biotin labelled target antibody by ELISA. Included as controls were the VHH72-Fc (17) and H11-H4-Fc (14) nanobodies linked to the hinge and Fc region of human IgG1, CR3022 and S309 human MAbs (11, 19) reconstituted as an IgG1 antibody. These four control molecules have characterised binding footprints on the RBD defined by crystal structures (12, 14, 17, 19), as does EY 6A (15). Also included was the protease domain (residues 18-615) of ACE2 linked to the Fc region of human IgG1 (ACE2-Fc dimer).

The ten antibodies formed four cross-inhibiting clusters (Table 2), represented by antibodies EY 6A (cluster 1, which included CR3022), FI 3A (cluster 2, which included H11-H4), FD 11A (cluster 3, which included S309) and FJ 10B (cluster 4). The strongest inhibitors of ACE2-Fc binding were in clusters 2 and 3 (Tables 1 and 2). Neutralising antibodies were detected in clusters 1, 2 and 3, with the strongest antibodies FI 3A and FD 11A being in clusters 2 and 3 (Tables 1 and 2).

**Table 2.**
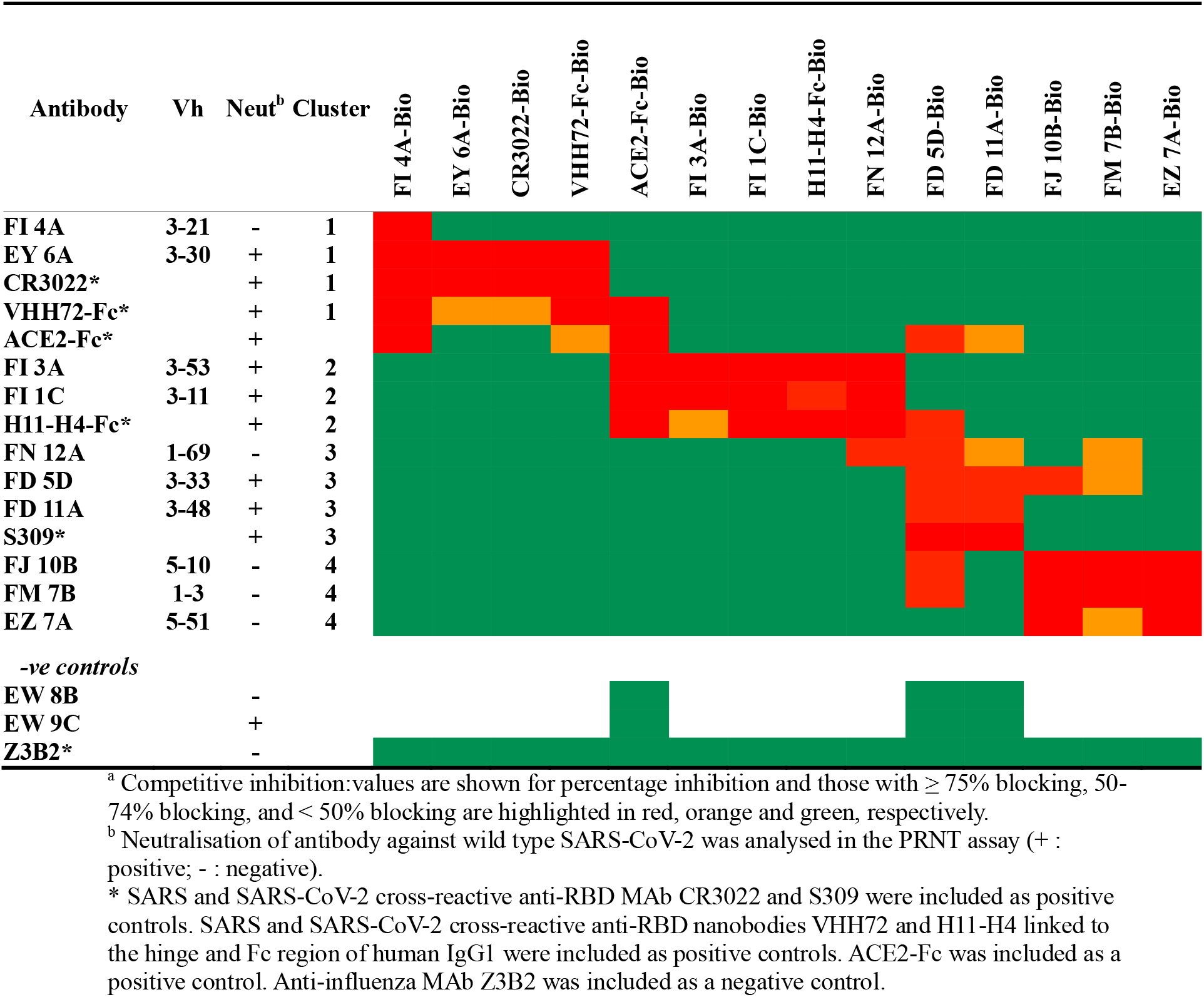
Competitive binding analysis of anti-SARS-CoV-2 RBD antibodies^a^

### Neutralising antibodies to RBD and relationship to ACE2 blockade

Five neutralising anti-RBD MAbs partially or completely blocked the interaction between RBD and ACE2 (Tables 1 and 2, Figure 3). The most potent neutralising antibodies were ACE2 blockers (FI 3A in cluster 2, and FD 11A in cluster 3), and bound independently of each other to the RBD (Figure 3c). MAb EY 6A has been shown to alter the binding kinetics of the interaction without full inhibition (15) and it had a moderate effect on ACE2 binding in the assay where ACE2 was expressed at the cell surface. These three MAbs bound independently of each other indicating the existence of at least three neutralisation-sensitive epitopes within the RBD (Figure 3c). All five neutralising MAbs to the RBD (EY 6A, FI 3A, FI 1C, FD 11A, FD 5A) had V gene sequences close to germline (Supplemental Table 2), indicating the development of *de novo* anti-RBD antibodies upon natural SARS-CoV-2 infection in humans.

The ability of convalescent sera to block soluble RBD for binding to cell-expressed ACE2 was assessed. Both convalescent sera from case A (D22) and case B (D26) exhibited detectable ACE2-blocking activities (Figure 3d). This serological activity is compatible with the isolation of potent ACE2-blocking antibodies from both patients (Table 1). By contrast, the D18 serum from case C showed minimal ACE2-blocking activity. Both cases A and B had prolonged fever and the development of pneumonia during hospitalization (Supplemental Table 1), which suggests that the development of ACE2-blocking antibody response in the convalescent stage is likely associated with clinical severity after infection.

### Neutralising antibodies to S2

Six of nine MAbs specific for S2 showed moderate neutralisation in the PRNT assay (Table 1). The antibodies FB 1E, FJ 4E and EW 9C, are moderately neutralising (EC50 36-133.33 nM), cross-react on the spike glycoprotein from the common cold betacoronavirus OC43, and show sequence characteristics of memory cells with high numbers of somatic mutations. This indicates that memory B cells, likely primed by an endemic or epidemic betacoronavirus related to OC43, can give rise to antibodies that neutralise SARS-CoV-2, albeit modestly. The other three neutralising antibodies specific for S2, FD 10A, FG 7A and FM 1A were close to germline in sequence (Supplemental Table 2) and did not cross-react strongly with other betacoronaviruses (Table 1). FD 10A exhibits the most potent neutralising activity in the PRNT assay and also completely inhibits SARS-CoV-2-induced cytopathic effect (see methods) at 8.33 nM.

### Neutralising antibodies to non-RBD S1

Thirteen MAbs were defined that bound the S1 region and three, close to germline in sequence, were neutralising. FJ 1C showed strong neutralisation (EC_50_ 55.5 nM), whilst FD 11E (EC_50_ 70 nM) and FD 1E (EC_50_ 110 nM) were moderately neutralising (Table 1). We investigated the specificity of these antibodies using flow cytometry with MDCK-SIAT1 cells expressing the N-terminal domain (NTD) of spike glycoprotein linked to the transmembrane and cytoplasmic region of influenza haemagglutinin (see methods). FJ 1C (neutralising) and FD 7C bound strongly to this NTD construct defining them as NTD-specific (Table 1, Supplemental Figure 7).

### Characterisation of anti-nucleocapsid antibodies

A set of 35 anti-SARS-CoV-2 nucleocapsid MAbs were derived from circulating plasmablasts in the naturally infected subjects (Table 3, Supplemental Figure 8); 19 of these strongly cross-react with at least one of the betacoronaviruses tested, 17 with SARS CoV, 14 with OC43 virus, and 13 with MERS CoV in ELISA. This suggested extensive cross-reactivity of the antibody response to epitopes in nucleocapsids of betacoronaviruses following natural infection with SARS-CoV-2.

**Table 3.** The antigenic specificity and cross-reactivity of 35 anti-SARS-CoV-2 nucleocapsid antibodies derived from COVID-19 patients.

| Antibody | Case | SARS-CoV-2<sup>a</sup> | Cross-reactivity<sup>a</sup> |  |  | Antibody | Case | SARS-CoV-2<sup>a</sup> | Cross-reactivity<sup>a</sup> |  |  |
| --- | --- | --- | --- | --- | --- | --- | --- | --- | --- | --- | --- |
|  |  |  | SARS | MERS | OC43 |  |  |  | SARS | MERS | OC43 |
| EY 12B | A | 1.23 | 1.24 | 1.10 | 1.24 | EZ 9B | B | 1.28 | 0.66 | 0.72 | 0.91 |
| EY 9C | A | 1.42 | 1.12 | 1.07 | 1.39 | FD 9B | B | 1.11 | 0.66 | 0.74 | 0.47 |
| EY 5A | A | 1.40 | 1.30 | 1.32 | 1.32 | FD 3E | B | 1.14 | 0.69 | 0.75 | 0.81 |
| EY 12A | A | 1.34 | 1.37 | 1.39 | 1.45 | FD 5B | B | 1.06 | 0.28 | 0.58 | 0.42 |
| EY 2A | A | 1.28 | 1.13 | 0.31 | 0.33 | EZ 7B | B | 1.01 | 0.29 | 0.29 | 0.33 |
| EY 3B | A | 1.22 | 1.00 | 0.22 | 0.26 | EZ 4A | B | 1.03 | 0.37 | 0.36 | 0.45 |
| EY 8A | A | 0.65 | 0.63 | 0.66 | 0.57 | FJ 3C | B | 0.98 | 1.28 | 1.16 | 0.86 |
|  |  |  |  |  |  | FD 4B | B | 0.98 | 0.43 | 0.44 | 0.46 |
| EZ 8C | B | 1.34 | 1.11 | 1.07 | 1.44 | FD 4E | B | 0.93 | 0.53 | 0.63 | 0.65 |
| EZ 8B | B | 1.38 | 1.28 | 1.24 | 1.12 | FD 4C | B | 0.84 | 0.47 | 0.59 | 0.54 |
| FJ 6D | B | 1.30 | 1.30 | 1.24 | 1.05 | FD 8C | B | 0.84 | 0.46 | 0.47 | 0.56 |
| EW 1A | B | 1.37 | 1.36 | 1.31 | 1.28 | EZ 4C-1 | B | 0.60 | 0.47 | 0.26 | 0.33 |
| EW 9A | B | 1.45 | 1.35 | 1.24 | 1.23 | EZ 4C-2 | B | 0.71 | 0.59 | 0.58 | 0.34 |
| EW 5A | B | 1.37 | 1.12 | 1.04 | 1.30 | EW 10C | B | 0.56 | 0.62 | 0.57 | 0.59 |
| EZ 9A | B | 1.51 | 1.28 | 1.29 | 1.13 | EZ 11A | B | 0.53 | 0.46 | 0.46 | 0.23 |
| EW 4C | B | 1.24 | 1.23 | 1.24 | 1.31 |  |  |  |  |  |  |
| FD 6D | B | 1.73 | 1.19 | 0.12 | 0.21 | FB 9B | C | 1.27 | 1.28 | 0.13 | 0.21 |
| EZ 11C | B | 1.46 | 0.73 | 0.72 | 1.16 | FL 9B | C | 0.92 | 0.60 | 0.68 | 0.94 |
| EZ 9C | B | 1.53 | 0.87 | 0.87 | 1.27 |  |  |  |  |  |  |
| *Controls* |  |  |  |  |  |  |  |  |  |  |  |
| CR3022 |  | 0.07 | 0.06 | 0.07 | 0.17 |  |  |  |  |  |  |
| BS-1A |  | 0.08 | 0.07 | 0.07 | 0.15 |  |  |  |  |  |  |

<sup>a</sup> A sample (10 µg/ml) is considered positive when the measured extinction is at least 3 times the OD<sub>450</sub> value of the negative control in the ELISA. CR3022 is an anti-SARS RBD human MAb and BS-1A is an anti-influenza H3 human MAb. OD values of $\geq 1.00$ or 0.50-0.99 or $\leq 0.49$ are highlighted in deep orange, orange and light orange, respectively.

The 35 MAbs were evolved from 33 clonal groups defined by their heavy chain VDJ and light chain VJ rearrangements (Supplemental Table 3). Those cross-reactive with other betacoronaviruses carried considerably more amino acid substitutions than SARS-CoV-1-specific MAbs in their variable domains (CR heavy 22±15 versus specific heavy 9±11, p=0.008, two-tailed Mann-Whitney test; Figure 2d), supporting the interpretation that approximately 50% of the anti-nucleocapsid antibody response in these three patients was derived from memory B cells, presumably primed by betacoronaviruses endemic to humans and related to OC43.

## Discussion

We detected a robust and rapid plasmablast response encoding diverse anti-spike glycoprotein and anti-nucleocapsid antibody populations within 3 weeks of onset of illness in COVID-19 patients. The concomitant serologic response occurred as early as the first week after illness onset, with gradually increasing levels of SARS-CoV-2 spike glycoprotein- and RBD-binding IgG antibodies from the second to third week after symptom onset. The kinetics of plasmablast and virus-specific serologic responses was observed to vary between subjects. Consecutive samples from donor B showed a highly expanded plasmablast subset at the time of progression into pneumonia, followed by class-switching to an IgG-predominant plasmablast response until week three of illness. On the other hand, donor C presented with mild symptoms and produced an early plasmablast response with class-switching at the end of first week of symptom onset. Thevarajan et al demonstrated a similar kinetic of B cell response in a mild COVID-19 case, in whom the peripheral plasmablast response and the SARS-CoV-2-binding IgG antibodies peaked at day eight, soon after the disappearance of fever (5). An early class-switching phenotype of the humoral response was also noted within the first week of onset in paediatric patients who mainly experienced mild illness after SARS-CoV-2 infection, although the immunological basis for this phenotype is unclear (20). Both peak plasmablast frequency and serologic titre for spike glycoprotein were higher in donor B, who presented with severe symptoms, than in donor C. Similar observations on the serologic titre and clinical severity have been reported by others (7, 21).

In this study, a substantial subset of plasmablast-derived anti-S2 (five of nine, 56%) and anti-nucleocapsid antibodies (14 of 35, 40%) cross-reacted with human betacoronavirus OC43. Cross-reactivity among betacoronavirus es has also been reported in polyclonal sera (22, 23). Human coronavirus OC43, discovered in the 1960s, is one of the major betacoronaviruses that cause common colds in the community (24, 25), and severe respiratory infections in elderly and immunocompromised individuals (23, 26). Epidemiologic surveys reveal that OC43 infection can occur in early childhood, and that OC43 seropositivity reaches nearly 90% in adults (27, 28). The S2 component of SARS-CoV-2 spike glycoprotein shares 43~89% amino acid identity with SARS, MERS and OC43, and, similarly, the amino acid sequence of SARS-CoV-2 nucleocapsid protein of is 34~90% homologous to those of other betacoronaviruses (Supplemental Figures 9 and 10), suggesting the presence of conserved epitopes on these antigens.

The presence of pre-existing immune memory to betacoronavirus that cross-react with SARS-CoV-2 is supported by the accumulation of somatic mutations in the genes encoding cross-reactive antibodies isolated from COVID-19 patients (Figures 2c and 2d, Supplemental Tables 2 and 3). This situation is reminiscent of re-exposure to immunogenic epitopes shared by closely related viruses leading to induction of broadly cross-reactive antibodies in patients infected with influenza, dengue or Zika viruses (29–31).

The 32 MAbs that bound to the spike glycoprotein were systematically tested for neutralisation (summarised in Table 1). Results established that neutralising epitopes were present on the RBD, S1-NTD, S1-non NTD/RBD, and S2 regions of the spike glycoprotein. The range of neutralisation EC_50_ titres against live SARS-CoV-2 reported in the literature for human and murine monoclonal antibodies span at least three orders of magnitude, from ng/ml to μg/ml (32, 33). Our results reflect the range of values in *in vitro* potency. The relationship between the EC_50_ value in neutralisation assays and therapeutic potential is not established, although antibodies to the RBD with EC_50_ in the nanogram range have shown therapeutic activity in small animal models (34–36). The majority of the most strongly neutralising antibodies were close to germline in sequence, showing that full affinity maturation is not required in order to achieve SARS-CoV-2 virus neutralisation. We made a similar observation for antibodies induced by an Ebola vaccine (18), and this has been observed in other collections of monoclonal antibodies to SARS-CoV-2 (4, 33–40).

The RBDs of SARS and SARS-CoV-2 are known to contain neutralising epitopes (15, 32–42), and vaccines based on the RBD of SARS and SARS-CoV-2 induce strong neutralising antibodies that are protective in animal models (43–45). There is a tendency for the most potent neutralising antibodies to be those that block the binding of the RBD to its receptor ACE2 (32–42, 46–48). Previous structural studies demonstrated that potent antibodies with footprints overlapping that of ACE2 are composed of two major subgroups, one with representative Vh3-53 gene usage and short CDR3 binding to the RBD in an up conformation and the other binding RBDs in both up and down conformations and even adjacent RBDs (49, 50). Potent MAb FI 3A shares similar characteristics with the first subgroup described above. Occasional antibodies that do not occupy the ACE2 footprint or weakly block ACE2 binding, i.e. S309, CR3022 and EY6A, can be almost as potent, perhaps through spike trimer cross-linking or triggering a conformational change in the spike glycoprotein that renders it non-functional (4, 15, 19, 40).

We characterise ten MAbs targeting the RBD that can be arranged into four cross-inhibiting clusters (Table 2). Three of these MAb clusters (represented by EY 6A, FI 3A and FD 11A) demonstrate neutralisation of SARS-CoV-2 to some level, while those in the fourth (represented by FM 7B) did not neutralise. This suggests that three neutralising antibodies may be able to bind to the RBD simultaneously. The most potent neutralising MAbs fall in two clusters and interfere strongly with ACE2 binding, which may be exploited therapeutically. For instance, antibodies FI 3A in cluster 2 (EC_50_ 8.67 nM) and FD 11A in cluster 3 (EC_50_ 0.05 nM) could be combined to limit possible selection of neutralisation-resistant variants. This principle has been demonstrated in recent studies (36, 38, 51).

Neutralisation by human MAbs to the S1-NTD of SARS-CoV-2 has been described (35, 52), but their mechanism of action is not known. The range of neutralising EC_50_ titres (0.09-51.1 nM) was overlapping with those targeting the RBD, similar to our NTD-specific antibody FJ 1C. Cocktails of antibodies that include a representative to the NTD would further reduce the likelihood of selecting neutralisation-resistant viruses. A second strong binder to the NTD, FD 7C, was not neutralising. Structural comparisons of these two antibodies bound to spike glycoprotein may provide insight into the neutralising action of FJ 1C.

We detected a subset of six MAbs to the S2 region of the spike glycoprotein that neutralised moderately (EC_50_ 36-133 nM). Three of these were clearly derived from a memory population; showing significant accumulation of somatic mutations in the MAb encoding genes and cross-reactivity for the OC43 common cold virus spike glycoprotein (Figure 2c). Further investigation is required to ascertain whether such antibodies, that may be weak- or non-neutralising and cross-reactive with common cold viruses, are beneficial or detrimental with respect to COVID-19 disease.

Our results have significance for serologic tests employing the N and S2 antigens of SARS-CoV-2. Serologic surveys, some of which are based on these antigens, with sera from donors infected with SARS-COV-2 during the spring and summer months have shown very high specificity (as judged by comparing convalescent sera from COVID-19 patients versus sera collected in the pre-COVID-19 period)(www.gov.uk/government/publications/covid-19-laboratory-evaluations-of-serological-assays). However, if many individuals have memory B cells to S2 and N antigens of circulating betacoronaviruses that cross-react with SARS-CoV-2, concurrent winter infections with these viruses might erode the specificity of serologic tests for SARS-CoV-2 based on these two antigens. Cross-reactivity of antibodies targeting the S1 subunit is relatively lower, so using this antigen is more likely to give a true indication of SARS-CoV-2 specificity.

In summary, COVID-19 patients developed strong anti-SARS-CoV-2 spike glycoprotein and nucleocapsid plasmablast and antibody responses. A panel of IgG MAbs targeted a diverse spectrum of epitopes on the RBD, S1-NTD, non-NTD/RBD S1 and S2 regions of the spike glycoprotein, of which 14 neutralised wild type virus with EC_50_s in the range 0.05 to ~133 nM. Neutralising activities of the majority of anti-RBD MAbs were linked to ACE2-binding blockade, and non-competing pairs of such MAbs, perhaps combined with a neutralising MAb to the NTD, offer potential formulations for the development of prophylactic and therapeutic agents against SARS-CoV-2. Antibody responses to nucleocapsid and the S2 component of spike glycoprotein confirm marked cross-reactivity with a common cold virus.

## Methods

### Study design

This study was designed to isolate SARS-CoV-2 antigen-specific MAbs from peripheral plasmablasts of humans infected with SARS-CoV-2 and to characterise the antigenic specificity and phenotypic activities of the MAbs. Diagnosis of SARS-CoV-2 infection was based on positive real-time reverse transcriptase polymerase chain reaction results of respiratory samples. The study protocol and informed consent were approved by the ethics committee at the Chang Gung Medical Foundation and the Taoyuan General Hospital, Ministry of Health and Welfare. Each patient provided signed informed consent. The study and all associated methods were carried out in accordance with the approved protocol, the Declaration of Helsinki and Good Clinical Practice guidelines.

### Staining and sorting of plasmablasts

Freshly separated peripheral blood mononuclear cells (PBMCs) or thawed PBMCs were stained with fluorescent-labelled antibodies to cell surface markers purchased from BD Biosciences, USA; Pacific blue anti-CD3 (clone UCHT1, Cat. No. 558117, BD), Fluorescein isothiocyanate anti-CD19 (clone HIB19, Cat. No. 555412, BD), Phycoerythrin-Cy7 anti-CD27 (clone M-T271, Cat. No. 560609, BD), Allophycocyanin-H7 anti-CD20 (clone L27, Cat. No. 641396, BD), Phycoerythrin-Cy5 anti-CD38 (clone HIT2, Cat. No. 555461, BD) and Phycoerythrin anti-human IgG (clone G18-145, Cat. No. 555787, BD). The CD3^neg^CD19^pos^CD20^neg^CD27^hi^CD38^hi^IgG^pos^ plasmablasts were gated and isolated in chamber as single cells as previously described (53).

### Production of human IgG 1 monoclonal antibodies

Sorted single cells were used to produce human IgG MAbs as previously described (53). Briefly, the variable region genes from each single cell were amplified in a reverse transcriptase polymerase chain reaction (RT-PCR: QIAGEN, Germany) using a cocktail of sense primers specific for the leader region and antisense primers to the Cγ constant region for heavy chain and Cκ and Cλ for light chain. The RT-PCR products were amplified in separate polymerase chain reactions for the individual heavy and light chain gene families using nested primers to incorporate restriction sites at the ends of the variable gene as previously described (53). These variable genes were then cloned into expression vectors for the heavy and light chains. Plasmids were transfected into the HEK293T cell line for expression of recombinant full-length human IgG MAbs in serum-free transfection medium. A selected panel of MAbs were further expanded and purified.

To determine the individual gene segments employed by VDJ and VJ rearrangements and the number of nucleotide mutations and amino acid replacements, the variable domain sequences were aligned with germline gene segments using the international ImMunoGeneTics (IMGT) alignment tool (http://www.imgt.org/IMGT_vquest/input).

### Enzyme-linked immunosorbent assay (ELISA)

ELISA plates (Corning® 96-well Clear Polystyrene High Bind Stripwell™ Microplate, USA) were coated with 8 μg/ml SARS-CoV-2 antigens (spike glycoprotein extracellular or receptor-binding domains, or nucleocapsid: Sino Biological, China) or SARS antigen (spike glycoprotein S1 subunit: Sino Biological, China) or Middle East Respiratory Syndrome coronavirus (MERS) antigen (spike glycoprotein extracellular domain: Sino Biological, China) or human coronavirus OC43 antigen (spike glycoprotein extracellular domain: Sino Biological, China) at 4°C overnight. Plates were washed with phosphate-buffered saline containing 0.05% Tween-20 and blocked with 3% bovine serum albumin (BSA) at room temperature for 1 hour on a shaker. Serial dilutions of MAb-containing cell culture supernatant or purified MAb were added and plates were incubated at 37°C for 1 hour. Plates were washed and incubated with horseradish peroxidase-conjugated rabbit anti-human IgG secondary antibody (Rockland Immunochemicals, USA). Plates were washed and developed with TMB substrate reagent (BD Biosciences, USA). Reactions were stopped with 0.5M hydrochloric acid and absorbances was measured at 450nm on a microplate reader. Non-transfected cell culture supernatant, anti-influenza H3 human IgG MAb BS 1A (in house), anti-SARS spike glycoprotein MAb CR3022 and convalescent serum were used as controls for each experiment. Reaction yielding an absorbance value above three times the mean absorbance of the negative control BS 1A were considered positive.

### Flow-cytometry based binding assay

MDCK-Spike cells were produced by stably transfecting parental MDCK-SIAT1 cells with cDNA expressing full-length SARS-CoV-2 spike glycoprotein. MDCK-RBD cells were produced by, stably transducing MDCK-SIAT1 cells with a Lentiviral vector encoding a cDNA expressing RBD amino acids 340-538 (NITN.GPKK underlined) fused via a short linker to the transmembrane domain of haemagglutinin H7 (A/Hong Kong/125/2017) (EPI977395) at the C-terminus for surface expression (sequence: MNTQILVFALIAIIPTNA/DKIGSGS<u>NITNLCPFGEVFNATRFASVYAWNRKRISN CVADYSVLYNSASFSTFKCYGVSPTKLNDLCFTNVYADSFVIRGDEVRQIAPGQTGKIADYNYKLPDDFTGCVIAWNSNNLDSKVGGNYNYLYRLFRKSNLKPFE RDISTEIYQAGSTPCNGVEGFNCYFPLQSYGFQPTNGVGYQPYRVVVLSFELL HAPATVCGPKK</u>TGSGGSGKLSSGYKDVILWFSFGASCFILLAIVMGLVFICVKN GNMRCTICI*). MDCK-NTD cells were produced by stably transfecting parental MDCK-SIAT1 cells with cDNA expressing SARS-CoV-2 NTD.

Both MDCK-Spike and MDCK-RBD cells were then FACS sorted for highly expressing cells using the CR3022 antibody. MDCK-Spike or MDCK-RBD cells were prepared and resuspended. Cells were probed with purified MAbs in 3% BSA. Bound primary antibodies were detected with FITC-conjugated anti-IgG secondary. The binding activities were analyzed by BD FACSCanto™ II flow cytometer (BD Biosciences, USA).

### Immunofluorescence assay

Under biosafety level 3 (BSL-3) conditions, Vero E6 cells were infected with 100 TCID_50_ (median tissue culture infectious dose) SARS-CoV-2 (hCoV-19/Taiwan/CGMH-CGU-01/2020, EPI_ISL_411915). Infected cells were placed on coverslips and, and fixed with acetone at room temperature for 10 minutes. After blocking with 1% BSA at room temperature for 1 hour and washing, fixed cells were incubated with MAb-containing cell culture supernatant. The anti-influenza human monoclonal antibody BS 1A, anti-SARS spike glycoprotein MAb CR3022 and convalescent serum were used as antibody controls for each experiment. Following incubation and wash, cells were stained with FITC-conjugated anti-human IgG secondary antibody and Evans blue dye as counterstain. Antibody-bound infected cells demonstrated an apple-green fluorescence against a background of red fluorescing material stained by the Evans Blue counterstain. Images were acquired with original magnification 40x, scale bar 20 μm.

### Plaque reduction neutralisation assay (Francis Crick Institute)

Confluent monolayers of Vero E6 cells in 96-well plates were incubated with ~14 plaque forming units (PFU) of SARS CoV-2 (hCoV-19/England/02/2020, EPI_ISL_407073) and antibodies in a 2-fold dilution series (triplicates) for 3 hours at room temperature. Inoculum was then removed, and cells were overlaid with plaque assay overlay. Cells were incubated at 37°C, 5% CO_2_ for 24 hours prior to fixation with 4% paraformaldehyde at 4°C for 30 minutes. Fixed cells were then permeabilised with 0.2% Triton-X-100 and stained with a horseradish peroxidase conjugated-antibody against virus protein for 1 hour at room temperature. TMB substrate was then added to visualise virus plaques as described previously for influenza virus (54). Convalescent serum from COVID-19 patients was used as a control.

### Fluorescent focus-forming units microneutralisation assay (FMNT) (Oxford)

In brief, this rapid, high-throughput assay determines the concentration of antibody that produces a 50% reduction in infectious focus-forming units of authentic SARS-CoV-2 in Vero cells, as follows. Triplicate serial dilutions of antibody are pre-incubated with a fixed dose of SARS-CoV-2 (Australia/VIC01/2020, GenBank MT007544) (55) in triplicate before incubation with Vero cells. A carboxymethyl cellulose-containing overlay is used to prevent satellite focus formation. Twenty hours post-infection, the monolayers are fixed with paraformaldehyde and stained for N antigen using MAb EY 2A. After development with a peroxidase-conjugated antibody and substrate, foci are enumerated by enzyme-linked immune absorbent spot reader. Data are analysed using four-parameter logistic regression (Hill equation) in GraphPad Prism 8.3.

### Quantitative PCR-based neutralisation assay

Neutralisation activity of MAb-containing supernatant was measured using SARS-CoV-2 (hCoV-19/Taiwan/CGMH-CGU-01/2020, EPI_ISL_411915) infected Vero E6 cells. Briefly, Vero E6 cells were pre-seeded in a 96 well plate at a concentration of 10^4^ cells per well. The following day, MAb-containing supernatants were mixed with equal volumes of 100 TCID_50_ virus preparation and incubated at 37°C for 1 hour, then mixtures were added to seeded Vero E6 cells and incubated at 37°C for 5 days. Cell, virus and virus back-titration controls were setup for each experiment. At day 5 the culture supernatant was harvested from each well, and virus RNA was extracted and quantified by real-time RT-PCR targeting the E gene of SARS-CoV-2 as previously described. The cycle threshold values of real-time RT-PCR were used as indicators of the copy number of SARS-CoV-2 RNA in samples with lower cycle threshold values corresponding to higher virus copy numbers.

### CPE-based neutralisation assay

Vero E6 cells in Dulbecco’s Modified Eagle’s Medium containing 10% FBS were added into 96-well plates and incubated at 37°C with 5% CO_2_ overnight to reach confluence. After washing with virus growth medium (VGM: Dulbecco’s Modified Eagle’s Medium containing 2% FBS), two-fold serially diluted MAbs in VGM starting at 100 μg/ml were added to each duplicated well. The plates were immediately transferred to a BSL-3 laboratory and 100 TCID_50_ SARS-CoV-2 (hCoV-19/Taiwan/4/2020, EPI_ISL_411927) in VGM was added. The plates were further incubated at 37°C with 5% CO_2_ for three days and the cytopathic morphology of the cells was recorded using an ImageXpress Nano Automated Cellular Imaging System.

### Competitive binding assays

Competitive binding assays were performed as described previously (18) with slight modifications for epitope mapping of the anti-RBD MAbs. Briefly, 0.5 μg/ml of RBD-VLP were coated on NUNC plates (50 μl per well) overnight at 4°C, washed and blocked with 300 μl of 5% (w/v) dried skimmed milk in PBS for 1 hour at room temperature prior to the assays. Antibody was biotinylated using EZ-Link Sulfo-NHS-LC-biotin (21237; Life Technologies) and then mixed with competing MAb (in at least 10-fold excess) and transferred to the blocked NUNC plates for 1 hour. A second layer Streptavidin-HRP (S911, Life Technologies) diluted 1:1,600 in PBS/0.1% BSA (37525; Thermo Fisher Scientific) was then added and incubated for another 1 hour. Plates were then washed, and signal was developed by adding POD substrate (11484281001, Roche) for 5 minutes before stopping the reaction with 1 M H_2_SO_4_. Absorbance (OD_450_) was measured using a Clariostar plate reader (BMG, Labtech). Mean and 95% confidence interval of 4 replicate measurements were calculated. Competition was measured as: (X-minimum binding/(maximum binding-minimum binding), where X is the binding of the biotinylated MAb in the presence of competing MAb. Minimum binding is the self-blocking of the biotinylated MAb or background binding. Maximum binding is binding of biotinylated MAb in the presence of non-competing MAb (anti-influenza N1 neuraminidase MAb).

### ACE2 blocking assays

Two assays were used to determine the blocking of binding of ACE2 to RBD by MAbs. RBD was anchored on the plate in the first assay whereas ACE2 was anchored for the second assay.

In the first ACE2 blocking assay, RBD-VLP (Spycatcher-mi3 VLP-particles conjugated with Spytagged-RBD recombinant protein) (16) was coated on ELISA plates as described for the competitive binding assay. Recombinant ACE2-Fc (18-615) protein expressed in Expi293F (Life Technologies) cells was chemically biotinylated using EZ-link Sulfo-NHS-Biotin (A39256; Life Technologies) and buffer exchanged to PBS using a Zebaspin desalting column (Thermo Fischer). MAbs were titrated in duplicate or triplicate as half-log serial dilution, 8-point series starting at 1 μM in 30 μl volume with PBS/0.1% BSA buffer. 30 μl of biotinylated ACE2-Fc at approx. 0.2 nM (40 ng/ml) was added to the antibodies. 50 μl of the mixture was transferred to the PBS-washed RBD-VLP coated plates and incubated for 1 hour at room temperature. Secondary Streptavidin-HRP antibody (S911, Life Technologies) diluted to 1:1600 was then added to the PBS-washed plates and incubated for 1 h at room temperature. Plates were then washed four times with PBS and signal was developed by adding POD substrate (11484281001, Roche) for 5 minutes before stopping with 1 M H_2_SO_4_. OD_450_ was measured using a Clariostar plate reader (BMG, Labtech). The control antibody (a non-blocking anti-influenza N1 MAb) or ACE2-Fc without antibody used to obtain the maximum signal and wells with PBS/BSA buffer only were used to determine the minimum signal. Graphs were plotted as % binding of biotinylated ACE2 to RBD. Binding % = {(X - Min)/(Max - Min)}*100 where X = measurement of the antibody, Min = buffer only, Max = biotinylated ACE2-Fc alone. 50% inhibitory concentrations of the antibodies against ACE2 was determined using non-linear regression curve fit using GraphPad Prism 8.

The second ACE2 blocking assay was performed as described previously (14, 15). Briefly, MDCK-SIAT1 cells were stably transfected to overexpress codon-optimised human ACE2 cDNA (NM_021804.1) using lentiviral vector and FACS sorted (MDCK-ACE2). Cells (3 x 10^4^ per well) were seeded on a flat-bottomed 96-well plate the day before the assay. RBD-6H (340-538; NITN.GPKK) was chemically biotinylated using EZ-link Sulfo-NHS-Biotin (A39256; Life Technologies). Serial half-log dilutions (starting at 1 μM) of antibodies and controls were performed in a U-bottomed 96 well plate in 30 μl volume. 30 μl of biotinylated RBD (25 nM) were mixed and 50 μl of the mixture was then transferred to the MDCK-ACE2 cells. After 1 hour a second layer Streptavidin-HRP antibody (S911, Life Technologies) diluted 1:1,600 in PBS/0.1% BSA (37525; Thermo Fisher Scientific) was added and incubated for another 1 hour. Plates were then washed four times with PBS and signal was developed by adding POD substrate (11484281001, Roche) before stopping with 1 M H_2_SO_4_ after 5 minutes. OD_450_ was measured using a Clariostar plate reader (BMG, Labtech). The control antibody (a non-blocking anti-influenza N1 antibody) was used to obtain maximum signal and PBS only wells were used to determine background. Graphs were plotted as % binding of biotinylated RBD to ACE2. The 50% inhibitory concentration of the blocking antibody was determined as described above.

## Data availability

Antibodies are available (by contacting Kuan-Ying A. Huang from the Chang Gung Memorial Hospital and Chang Gung University and Alain R. Townsend from Oxford University) for research purposes only under an MTA, which allows the use of the antibody for non-commercial purposes but not their disclosure to third parties. The data that support the findings of this study are available from the corresponding authors on request.

## Author contributions

K.-Y.A.H. conceived and designed the study of MAb isolation and characterisation. K.-Y.A.H. produced and characterised plasmablast-derived MAbs. A.R.T. conceived and designed the study of MAb characterisation at Oxford. P.R, T.K.T, L.S generated cell lines, expressed proteins and antibodies, and performed experiments. S.H., R.H. R.S.D and J.W.M. designed and performed neutralisation assay at Crick Institute. A. H., J. G-J., X. L., M. K. and W.J. designed and performed neutralisation assay at Oxford. T-H.C., C-G.H., C-P.C., S-R.S, Y-C.L., C-Y.C., S-H.C., Y-C.H., T-Y.L., J-T.J. and C.M helped prepare materials, perform experiments and analyse data. All authors read and approved the manuscript.

## Acknowledgements

We acknowledge the BD FACSAria™ cell sorter service provided by the Core Instrument Center of Chang Gung University. Plasmablast sorting, production and characterisation of human MAbs were supported by the Chang Gung Memorial Hospital (BMRPE22). We would like to acknowledge Paul Sopp and Craig Waugh in the flow cytometry facility at the MRC WIMM for providing cell sorting services. The facility is supported by the MRC HIU, MRC MHU (CC_UU_12009); NIHR Oxford BRC; Kay Kendall Leukaemia Fund (KKL1057), John Fell Fund (131/030 and 101/517), the EPA fund (CF182 and CF170) and by the MRC WIMM Strategic Alliance awards G0902418 and MC_UU_12025. Production of antibodies was funded by the Fast Grant Application given to A.R.T., P.R., and T.K.T. P.R., L.S. and A.R.T. are funded by the Chinese Academy of Medical Sciences (CAMS) Innovation Fund for Medical Science (CIFMS), China (grant no. 2018-I2M-2-002). T.K.T. is funded by the EPA Cephalosporin Fund and The Townsend–Jeantet Charitable Trust (charity no. 1011770). The work done at the Crick Worldwide Influenza Centre was supported by the Francis Crick Institute receiving core funding from Cancer Research UK (FC001030), the Medical Research Council (FC001030) and the Wellcome Trust (FC001030). The Oxford work was funded in part through the generous support of philanthropic donors to the University of Oxford’s COVID-19 Research Response Fund.

