## Supplemental for "Breadth and function of antibody response to acute SARS-CoV-2 infection in humans"

| **Supplemental Table 1. Clinical characteristics of COVID-19 patients and sampling dates in the study.** | | | | | | | | | | | |
| --- | --- | --- | --- | --- | --- | --- | --- | --- | --- | --- | --- |
|  | **Age (yrs)** | | **Gender** | **Onset symptoms** | **Pneumonia** | **Oxygen  use** | **Ventilator  use** | **Fever subside** | **Sequela** | **Plasmablast sampling dates** | **Serum sampling dates** |
| **Case A** | 43 | M | | Fever, headache | Day 7  after onset | Yes | none | Day 9 after onset | none | Day 14  after onset | Day 22  after onset |
| **Case B** | 55 | F | | Cough, fever | Day 11  after onset | Yes | none | Day 20 after onset | none | Days 14, 18 and 22  after onset | Days 14, 18 and 26  after onset |
| **Case C** | 52 | M | | Fever | none | none | none | Day 1 after onset | none | Days 2, 6 and 14  after onset | Days 6, 10, 14 and 18  after onset |

| **Supplemental Table 2. Anti-SARS-CoV-2 spike monoclonal antibody heavy and light chain variable domain gene usage.** | | | | | | | | | | | | | |
| --- | --- | --- | --- | --- | --- | --- | --- | --- | --- | --- | --- | --- | --- |
| MAb | H-L | Vh | Jh | Dh | rf | V_H_ junction sequence | nt Mut | aa Sub | Vl | Jl | V_L_ Junction Sequence | nt Mut | aa Sub |
| FM 7B | H-𝝺 | 1-3*04 F | 5*02 F | 2-2*01 F | 2 | CARDPTYCSSTSCYPFSWFDPW | 3 | 1 | 3-21*02 F | 3*02 F | CQVWDSTGDHSWVF | 2 | 2 |
| FD 8B | H-K | 1-24*01 F | 6*02 F | 2-2*01 F | 2 | CATAAAINCSSTSCYYYYYYYGMDVW | 0 | 0 | 2-24*01 F | 2*01 F | CTQATQFPYTF | 2 | 2 |
| EW 9B | H-K | 1-46*01 or 03 F | 6*02 F | 2-2*01 F | 3 | CAREDGVVPAANLMISLEDYYYYGMDVW | 2 | 2 | 3-11*01 F | 4*01 F | CQQRSNWPLTF | 0 | 0 |
| FN 12A | H-𝝺 | 1-69*04 or 09 F | 6*02 F | 2-2*01 F | 2 | CARSGCSSTSCPSNLYYYYYGMDVW | 1 | 0 | 1-51*01 F | 3*02 F | CGTWDSSLSALVF | 1 | 0 |
| FG 12C | H-𝝺 | 3-9*01 F | 4*02 F | 4-17*01 F | 2 | CAKDMRVHDYGDYYFDYW | 0 | 0 | 3-21*01 F | 2*01 or 3*01 F | CQVWDSSSDHPVF | 1 | 1 |
| **FI 1C** | H-𝝺 | 3-11*04 F | 3*02 F | 6-13*01 F | 1 | CARRSNRFLIAFDIW | 3 | 2 | 2-14*01 F | 2*01 or 3*01 F | CSSYTSSSTLVVF | 1 | 1 |
| FI 4A | H-𝝺 | 3-21*01 F | 4*02 F | 2-21*02 F | 2 | CATYLFGDSHTYW | 9 | 7 | 6-57*02 F | 3*02 F | CQSYDSSNLHWVF | 0 | 0 |
| FD 1E | H-𝝺 | 3-21*01 F | 6*02 F | 6-13*01 F | 2 | CASLAAAGPETYYYYGMDVW | 1 | 0 | 3-1*01 F | 2*01 or 3*01 F | CQAWDSSVVF | 0 | 0 |
| FD 11E | H-𝝺 | 3-21*01 F | 6*02 F | 6-13*01 F | 2 | CASLAAAGPETYYYYGMDVW | 0 | 0 | 3-1*01 F | 2*01 or 3*01 F | CQAWDSSVVF | 0 | 0 |
| FN 2C | H-𝝺 | 3-30*03 or 18 or 3-30-5*01 F | 3*01 or 02 F | 3-10*01 F | 1 | CAKRREIFWLGEPPLSDAFDFW | 22 | 14 | 1-40*01 F | 3*02 F | CQSYDSSLSGSVF | 8 | 4 |
| **EY 6A** | H-K | 3-30*03 or 18 or 3-30-5*01 F | 4*02 F | 2-21*01 F | 1 | CAKDGGKLWVYYFDYW | 6 | 5 | 1-39*01 F or 1D-39*01 F | 4*01 F | CQQSYSTLALTF | 0 | 0 |
| FD 11D | H-K | 3-30*03 or 18 or 3-30-5*01 F | 4*02 F | 6-19*01 F | 1 | CAKEGAGSGWYRHHKPGYYFDYW | 1 | 1 | 3-20*01 F | 1*01 F | CQQYGSSPLTF | 0 | 0 |
| EW 9C | H-K | 3-30*03 or 18 or 3-30-5*01 F | 5*01 or 02 F | 3-10*01 F | 1 | CARATSIFWFGEGRNWFDPW | 33 | 15 | 3-11*01 F | 4*01 F | CQQRSNWPLTF | 25 | 13 |
| FD 1D | H-K | 3-30*04 or 3-30-3*03 F | 5*02 F | 3-10*01 F | 2 | CARAGSGSYLNWFDPW | 1 | 0 | 3-11*01 F | 5*01 F | CQQRSNWPITF | 0 | 0 |
| FG 7A | H-K | 3-30-3*01 F | 4*02 F | 1-26*01 F | 3 | CARSHSGSYRASLDYW | 2 | 2 | 3-20*01 F | 2*01 F | CQQYGSSPLYTF | 0 | 0 |
| FM 1A | H-𝝺 | 3-33*01 or 06 F | 4*02 F | 1-26*01 F | 1 | CAREGAVGATRGFDYW | 1 | 0 | 3-21*02 F | 2*01 or 3*01 F | CQVWDSSSDQGVF | 2 | 1 |
| **FD 11A** | H-𝝺 | 3-33*01 or 06 F | 6*02 F | 3-9*01 F | 2 | CAKGPDILTGYYNYYYYGMDVW | 2 | 2 | 1-40*01 F | 2*01 or 3*01 F | CQSYDSSLSGFYVVF | 0 | 0 |
| FN 8C | H-𝝺 | 3-33*05 F | 6*02 F | 3-9*01 F | 2 | CARERTYYDILTGYRHYYGMDVW | 1 | 1 | 3-21*02 F | 3*02 F | CQVWDSSSDHWVF | 1 | 1 |
| FD 5E | H-K | 3-43D*03 F | 6*02 F | 3-3*01 F | 1 | CAKDSVRFRYYYGMDVW | 0 | 0 | 3-11*01 F | 3*01 F | CQQRSNWPLTF | 0 | 0 |
| **FD 5D** | H-K | 3-48*04 F | 6*02 F | 6-13*01 F | 2 | CASPGGITAAGTSVLFGYYGMDVW | 4 | 2 | 2-28*01 or 2D-28*01 F | 1*01 F | CMQALQTPITWTF | 0 | 0 |
| **FI 3A** | H-K | 3-53*01 F | 3*02 F | 6-6*01 F | 3 | CARDHVRPGMNIW | 2 | 2 | 1-33*01 or 1D-33*01 F | 4*01 F | CQQYDNLPVTF | 1 | 0 |
| **FD 10A** | H-K | 3-74*01 F | 3*02 F | - |  | CANMAFDIW | 0 | 0 | 4-1*01 F | 5*01 F | CQQYYSTPITF | 0 | 0 |
| FJ 4E | H-K | 4-31*06 F | 5*02 F | 3-10*01 F | 2 | CARDEYDSSDSGIQGHWFDPW | 20 | 15 | 1-39*01 F or 1D-39*01 F | 1*01 F | CQQSYSTPWTF | 15 | 10 |
| **FJ 1C** | H-𝝺 | 4-38-2*02 F | 4*02 F | 3-10*01 F | 1 | CARDKALLWFGELFTNLFDYW | 0 | 0 | 2-14*01 F | 2*01 or 3*01 F | CSSYTSSSTLVF | 1 | 1 |
| EW 8B | H-K | 4-39*01 F | 3*02 F | 3-16*01 F | 2 | CARQEVWGGFDIW | 20 | 12 | 3-20*01 F | 1*01 F | CQQYGSSPTF | 4 | 4 |
| FD 11C | H-𝝺 | 4-39*07 F | 6*02 F | 3-10*01 F | 2 | CAREYYYGSETKKYYYYYGMDVW | 2 | 1 | 3-1*01 F | 2*01 or 3*01 F | CQAWDSSTVF | 0 | 0 |
| FD 7C | H-K | 4-59*01 F | 5*02 F | 3-10*02 F | 3 | CARDYRFGELFGRFAWFDPW | 1 | 1 | 3-15*01 F | 1*01 F | CQQYNNWPRAF | 2 | 1 |
| FD 7D | H-𝝺 | 5-10-1*03 F | 3*02 F | 2-2*01 F | 2 | CARHSDCSSTSCYFVDAFDIW | 1 | 1 | 3-25*03 F | 2*01 or 3*01 F | CQSADSSGTYVVF | 1 | 0 |
| FJ 10B | H-K | 5-10-1*03 F | 6*02 F | 3-16*01 F | 1 | CARLDPRYGPDYYGMDVW | 2 | 1 | 1-39*01 F or 1D-39*01 F | 4*01 F | CQQSYSTPLTF | 3 | 2 |
| FB 9D | H-𝝺 | 5-51*01 F | 6*02 F | 3-10*01 F | 3 | CARHWASMVRGVIRASHYYGMDVW | 27 | 11 | 2-8*01 F | 3*02 F | CSSYAFGGSDTRVF | 20 | 10 |
| FB 1E | H-𝝺 | 5-51*01 F | 6*02 F | 3-10*01 F | 3 | CARHWASMVRGVIRASHYYGMDVW | 22 | 11 | 2-8*01 F | 3*02 F | CSSYAFGGSDIRVF | 16 | 8 |
| EZ 7A | H-𝝺 | 5-51*01 F | 6*02 F | 5-12*01 F | 3 | CARGWVYRGFPYYGMDVW | 2 | 1 | 2-11*01 F | 3*02 F | CCSYAGSYTLVF | 0 | 0 |

Abbreviations: H, heavy; K, kappa; 𝝺, lambda; Vh, variable gene segment of the heavy chain variable domain; Dh, diversity gene segment of the heavy chain variable domain; Jh, joining gene segment of the heavy chain variable domain; Mut, number of nucleotide mutations; Sub, number of amino acid substitutions; Vl, variable gene segment of the light chain variable domain; Jl, joining gene segment of the light chain variable domain.

| **Supplemental Table 3. Anti-SARS-CoV-2 nucleocapsid monoclonal antibody heavy and light chain variable domain gene usage.** | | | | | | | | | | | | | |
| --- | --- | --- | --- | --- | --- | --- | --- | --- | --- | --- | --- | --- | --- |
| MAb | H-L | VH | JH | DH | rf | V_H_ junction sequence | nt Mut | aa Sub | VL | JL | V_L_ Junction Sequence | nt Mut | aa Sub |
| EZ 9B | H-𝝺 | 1-2*06 F | 5*02 F | 4-17*01 F | 3 | CAREGPTVTWWFDPW | 0 | 0 | 3-21*02 F | 3*02 F | CQVWDSSSDHPSWVF | 0 | 0 |
| EZ 11C | H-𝝺 | 1-24*01 F | 5*02 F | 4-17*01 F | 3 | CATTTVTTPTANWFDPW | 0 | 0 | 1-51*02 F | 2*01 or 3*01 F | CGTWDSSLRQVVF | 3 | 1 |
| FD 9B | H-𝝺 | 3-7*01 F | 2*01 F | 2-21*01 F | 1 | CVKFGRSEGLFW | 21 | 17 | 7-46*01 F | 2*01 or 3*01 or 3*02 F | CFLTYVGARRLF | 6 | 4 |
| EZ 11A | H-𝝺 | 3-7*01 F | 4*02 F | 1-26*01 F | 3 | CARDDYSGSYYWEFDYW | 0 | 0 | 4-69*01 F | 3*02 F | CQTWGTGIWVF | 0 | 0 |
| EW 10C | H-K | 3-7*03 F | 4*02 F | 4-23*01 ORF | 1 | CARGRTLGDW | 25 | 15 | 2-30*02 F | 3*01 F | CMQGTHWPPITF | 3 | 1 |
| EY 12B | H-𝝺 | 3-9*01 F | 3*01 F | 5-12*01 F | 3 | CAKGRSGYGHTAFDVW | 16 | 11 | 1-40*01 or 02 F | 2*01 or 3*01 F | CQSYDSSLSASVF | 9 | 6 |
| EZ 8C | H-𝝺 | 3-21*01 F | 3*02 F | 5-18*01 F | 3 | CARELTSYGSHDAFDIW | 15 | 9 | 2-14*01 F | 2*01 or 3*01 F | CSSYTTTDSVVF | 11 | 6 |
| EY 9C | H-𝝺 | 3-21*01 F | 4*03 F | 2-21*01 F | 2 | CATWGGAPFDYW | 12 | 9 | 2-23*01 or 02 or 03 F | 2*01 or 3*01 F | CCSYAGGRTFNVLF | 12 | 10 |
| EZ 8B | H-𝝺 | 3-23*04 F | 5*01 or 02 F | 3-16*01 F | 2 | CAKDLGYYGSGSSW | 4 | 2 | 3-1*01 F | 3*02 F | CQAWDSSTAVF | 0 | 0 |
| FD 3E | H-K | 3-30*03 or 18 or 3-30-5*01 F | 6*02 F | 3-10*01 F | 2 | CAKDPHYYGSGSYYNQLRGYYYYGMDVW | 1 | 1 | 2-29*02 F | 4*01 F | CMQGIHLP#TF | 0 | 0 |
| EW 1A | H-𝝺 | 3-33*01 or 06 F | 3*01 F | 6-13*01 F | 3 | CARDGQHLAPFAMDVW | 32 | 17 | 2-14*01 F | 3*02 F | CNSFVSGDSWVF | 27 | 17 |
| FD 5B | H-𝝺 | 3-33*01 or 06 F | 4*02 F | 5-24*01 ORF | 3 | CARDERRESYNFVLDYW | 6 | 3 | 2-8*01 F | 1*01 F | CSSYAGSNNPFVF | 1 | 1 |
| EW 9A | H-𝝺 | 3-33*01 or 06 F | 6*02 F | 3-9*01 F | 2 | CAKDMWALYDILTGYYTPYYYYGMDVW | 1 | 1 | 3-25*03 F | 3*02 F | CQSADSSGTYWVF | 1 | 1 |
| FD 8C | H-K | 3-49*05 F | 4*02 F | 3-3*01 F | 2 | CTRNDFWSGYYPDYW | 0 | 0 | 2-30*02 F | 4*01 F | CMQGTHWP#LTF | 0 | 0 |
| EW 5A | H-𝝺 | 3-64*05 or 3-64D*06 F | 2*01 F | 3-3*01 F | 2 | CVKDRGSVIRDFDVW | 37 | 23 | 7-43*01 F | 2*01 or 3*01 or 3*02 F | CLLYCGGGQLF | 21 | 13 |
| EZ 9C | H-𝝺 | 3-64D*06 F | 4*02 F | 6-19*01 F | 1 | CGKGLLSASGGLPIDDW | 34 | 20 | 1-51*01 F | 3*02 F | CATWDSSLSAGVF | 5 | 3 |
| EZ 9A | H-𝝺 | 3-74*01 F | 4*02 F | 4-11*01 ORF | 2 | CARDVNRYPDYW | 29 | 18 | 2-14*01 F | 3*02 F | CCSYVNNGAWVF | 28 | 13 |
| EY 12A | H-𝝺 | 4-30-4*01 F | 2*01 F | 3-9*01 F | 2 | CARGMTQDDILTGFNRPHWYFDLW | 22 | 12 | 1-40*01 F | 2*01 or 3*01 F | CQSFDSSLSDFVVF | 8 | 6 |
| FD 4E | H-𝝺 | 4-31*03 F | 4*02 F | 1-26*01 F | 3 | CARGRGSYLAGGNYYFDYW | 0 | 0 | 6-57*01 F | 3*02 F | CQSYDSSN#VF | 2 | 2 |
| FD 4C | H-𝝺 | 4-31*03 F | 4*02 F | 6-13*01 F | 1 | CARVRSSSSWYFDYW | 2 | 1 | 2-14*01 F | 3*02 F | CSSYTSKWVF | 4 | 2 |
| EW 4C | H-𝝺 | 4-38-2*02 F | 1*01 F | 3-16*01 F | 2 | CVRGTYGSGLHW | 49 | 24 | 7-46*01 F | 3*02 F | CFLSHNDAWVF | 17 | 12 |
| FD 6D | H-K | 4-38-2*02 F | 3*01 or 02 F | 3-22*01 F | 2 | CARDRLLAVHYDSRGYLVDYW | 22 | 10 | 4-1*01 F | 1*01 F | CQQYYDIPRTF | 12 | 6 |
| EY 5A | H-𝝺 | 4-59*01 F | 4*02 F | 4-23*01 ORF | 3 | CARGPGPATGGSLDYW | 23 | 16 | 1-44*01 F | 7*01 F | CSAWDDSLNGPVF | 7 | 5 |
| EZ 7B | H-K | 4-59*13 F | 3*02 F | 1-26*01 F | 3 | CARRVFGPVLPSKLGGSYWGGGAFDIW | 1 | 1 | 4-1*01 F | 3*01 F | CQQYYSTPLTF | 0 | 0 |
| EZ 4C-1 | H-K | 4-61*01 or 03 F | 4*02 F | 3-3*01 F | 1 | CARAPSAPFGGLFDWILPKGINNW | 24 | 15 | 1-5*03 F | 2*03 F | CQQYNGYSYSF | 17 | 8 |
| EZ 4C-2 | H-𝝺 | 4-61*01 or 03 F | 4*02 F | 3-3*01 F | 1 | CARAPSAPFGGLFDWILPKGIDNW | 22 | 13 | 1-41*01 ORF | 3*02 F | C*IA*HSSPR#WVF (2nd-CYS 104 not identified) | 14 | 8 |
| FD 4B | H-𝝺 | 4-61*01 or 03 F | 5*02 F | 3-3*01 F | 1 | CARAPSAPFGGLFDWILPKGIDSW | 24 | 13 | 1-41*01 ORF | 3*02 F | C*IA*HSSPR#WVF (2nd-CYS 104 not identified) | 14 | 8 |
| EY 8A | H-𝝺 | 4-61*02 F | 6*02 F | 5-12*01 F | 3 | CAKGHVISGYDDYYYYYGMDVW | 1 | 1 | 3-25*03 F | 3*02 F | CQSADSSGTYWVF | 0 | 0 |
| EY 2A | H-K | 5-51*01 F | 4*02 F | 6-13*01 F | 1 | CVRQERGSNTWYAGNSW | 44 | 22 | 2-28*01 or 2D-28*01 F | 2*02 F | CMQALQTPGTF | 14 | 7 |
| EY 3B | H-K | 5-51*01 F | 4*02 F | 6-13*01 F | 1 | CVRQERGSNTWYAGNSW | 41 | 21 | 2-28*01 or 2D-28*01 F | 2*02 F | CMQALQTPGTF | 13 | 7 |
| EZ 4A | H-K | 5-51*01 F | 4*02 F | 6-13*01 F | 2 | CARSPIAADLFDYW | 0 | 0 | 1-33*01 or 1D-33*01 F | 3*01 F | CQQYDNLLFTF | 0 | 0 |
| EZ 8B | H-K | 3-23*04 F | 5*01 or 02 F | 3-16*01 F | 2 | CAKDLGYYGSGSSW | 4 | 2 | 3-7*04 ORF | 1*01 F | CQQDYNS#TF | 1 | 1 |

Abbreviations: H, heavy; K, kappa; 𝝺, lambda; Vh, variable gene segment of the heavy chain variable domain; Dh, diversity gene segment of the heavy chain variable domain; Jh, joining gene segment of the heavy chain variable domain; Mut, number of nucleotide mutations; Sub, number of amino acid substitutions; Vl, variable gene segment of the light chain variable domain; Jl, joining gene segment of the light chain variable domain.

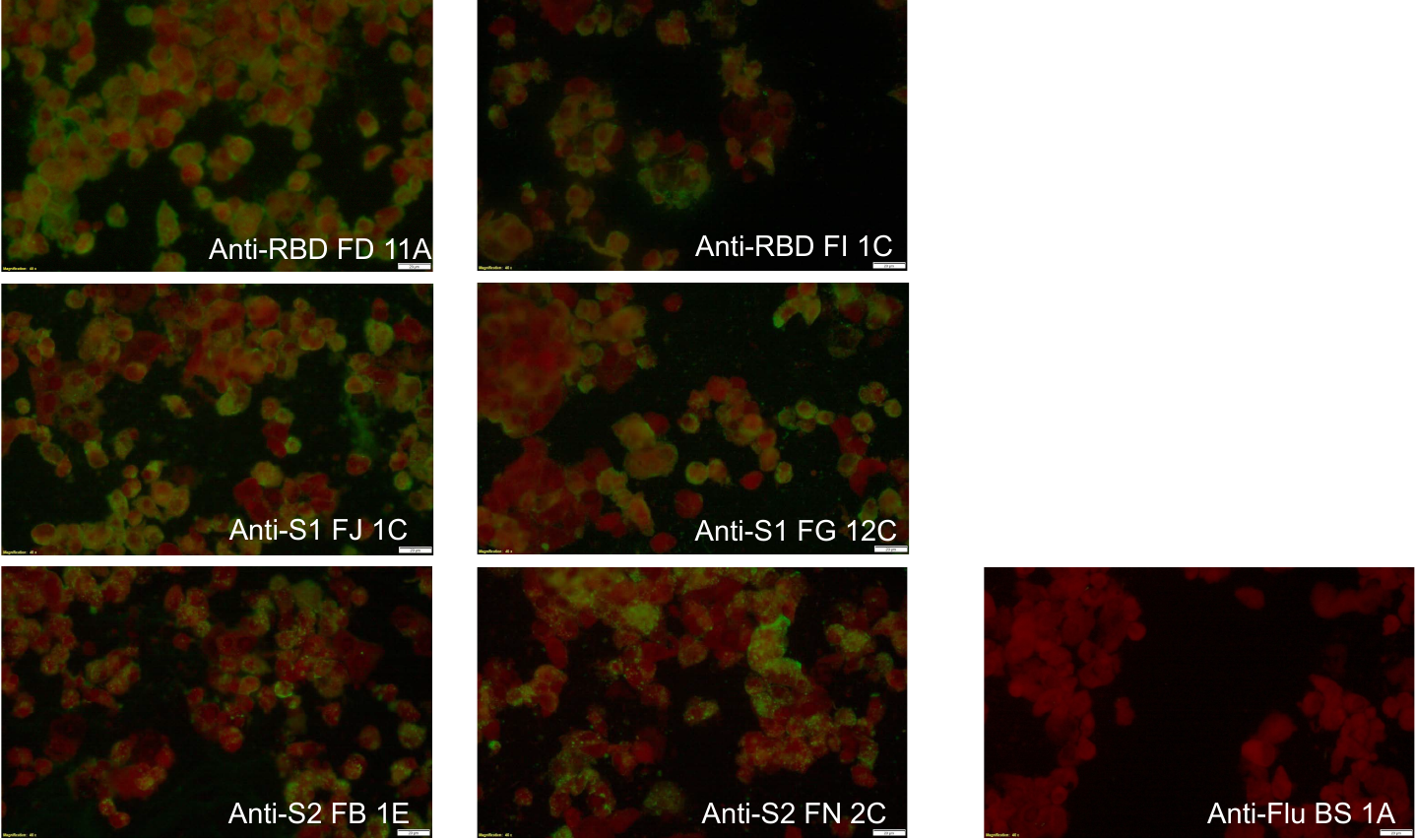

**Supplemental Figure 1. Binding of anti-spike antibodies to SARS-CoV-2-infected cells in immunofluorescence assay.** Representative immunofluorescence staining of anti-RBD, anti-S1 and anti-S2 MAbs are shown as apple-green fluorescence a background of red fluorescing material stained by Evans Blue counterstain. Anti-influenza H3 MAb BS 1A was included as a control. Images were acquired with original magnification 40x, scale bar 20 µm.

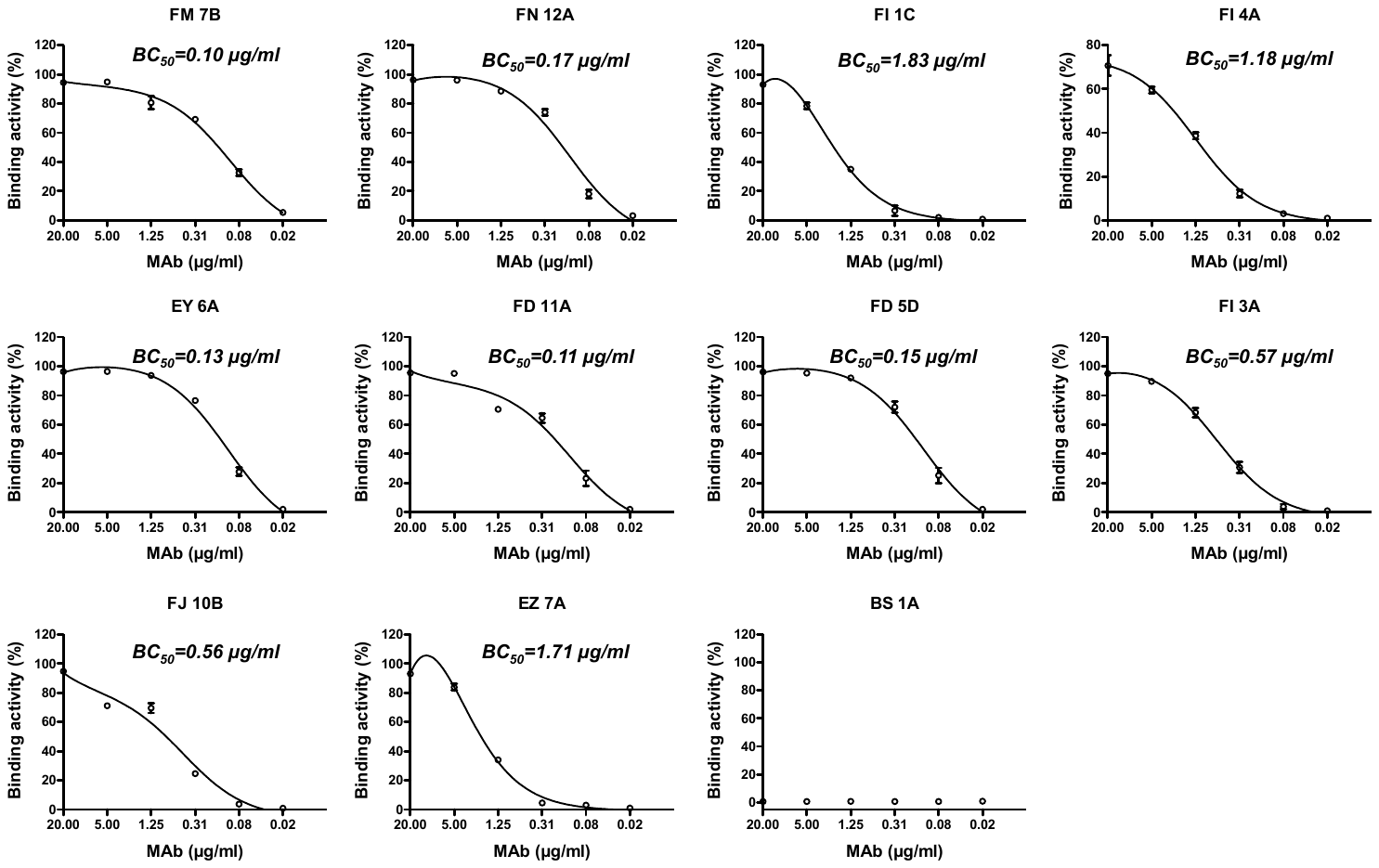

**Supplemental Figure 2. Binding activity of anti-SARS-CoV-2 RBD antibodies to MDCK-RBD cells measured by flow cytometry.** Anti-influenza H3 MAb BS-1A was included as a control. Binding percentages are presented as mean ± standard error of the mean. Each experiment was repeated twice (n=2). The 50% binding concentration (BC_50_) was measured with a curve fit using non-linear regression.

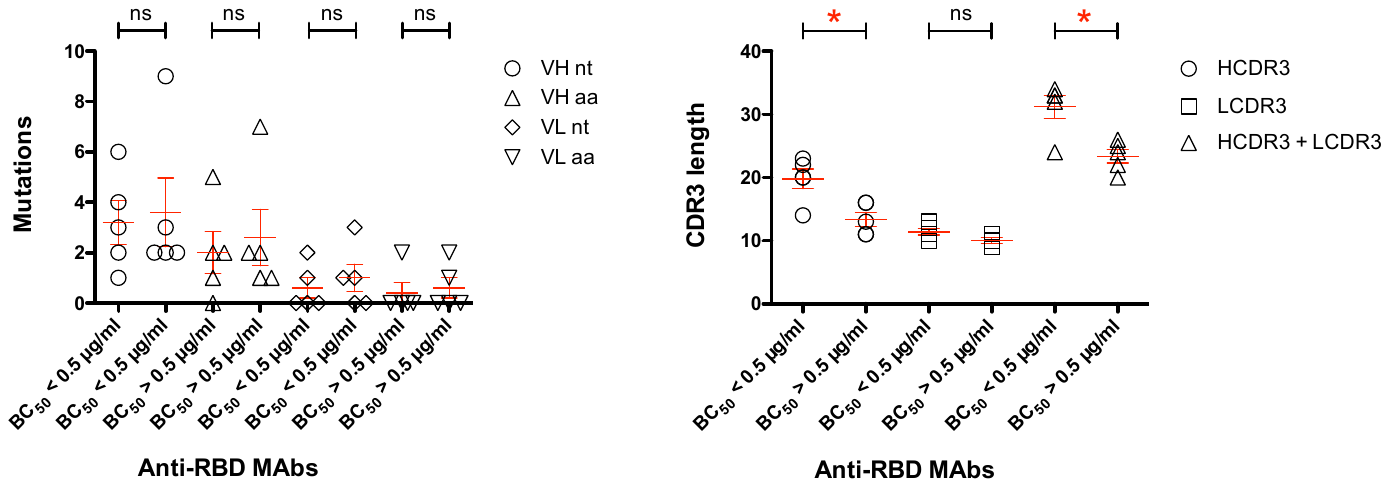

**Supplemental Figure 3. The correlation of CDR3 length with BC_50_ among anti-RBD antibodies.** The CDR3 length (number of amino acids) and MAb gene mutation numbers are presented as mean ± standard error of the mean (< 0.5 µg/ml, n=5 versus > 0.5 µg/ml, n=5). The two-tailed Mann-Whitney test was performed to compare the CDR3 length and mutation numbers between two groups. * P < 0.05 ; ns, non-significant ; BC_50_, 50% binding concentration.

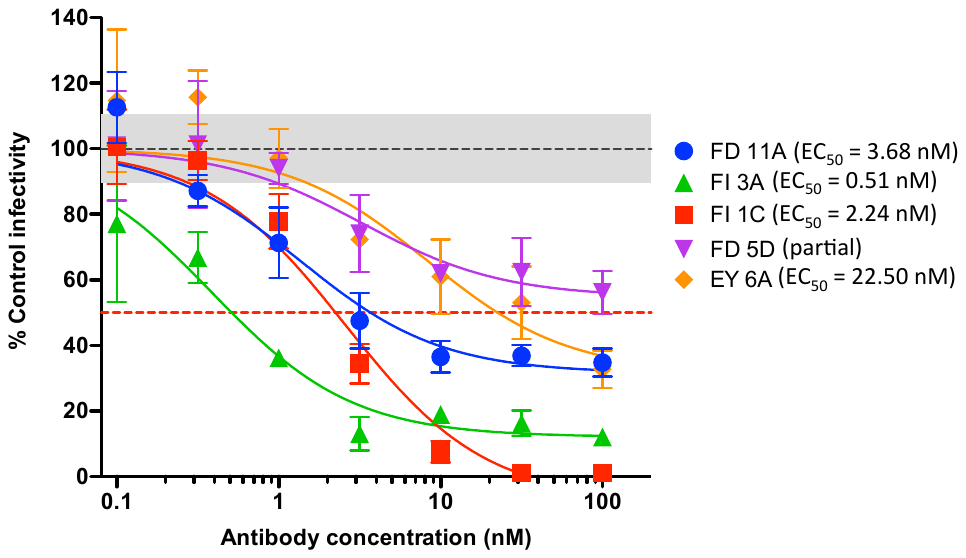

**Supplemental Figure 4. Neutralisation of wild type SARS-CoV-2 by anti-RBD monoclonal antibodies.** Neutralisation assays were performed on the indicated antibodies according to the fluorescent focus-forming units microneutralisation method (see methods). Data were normalized to control (no antibody) values of foci, and the grey region comprises ± 1 standard deviation the mean control values. Individual points are displayed ± 1 standard deviation of technical, and curves are shown only where the data for a particular antibody fitted the standard dose-response (Hill) equation (n=3). Partial: MAb neutralises at least ~40% viruses at 100 nM (highest concentration tested). EC_50_, 50% effective concentration.

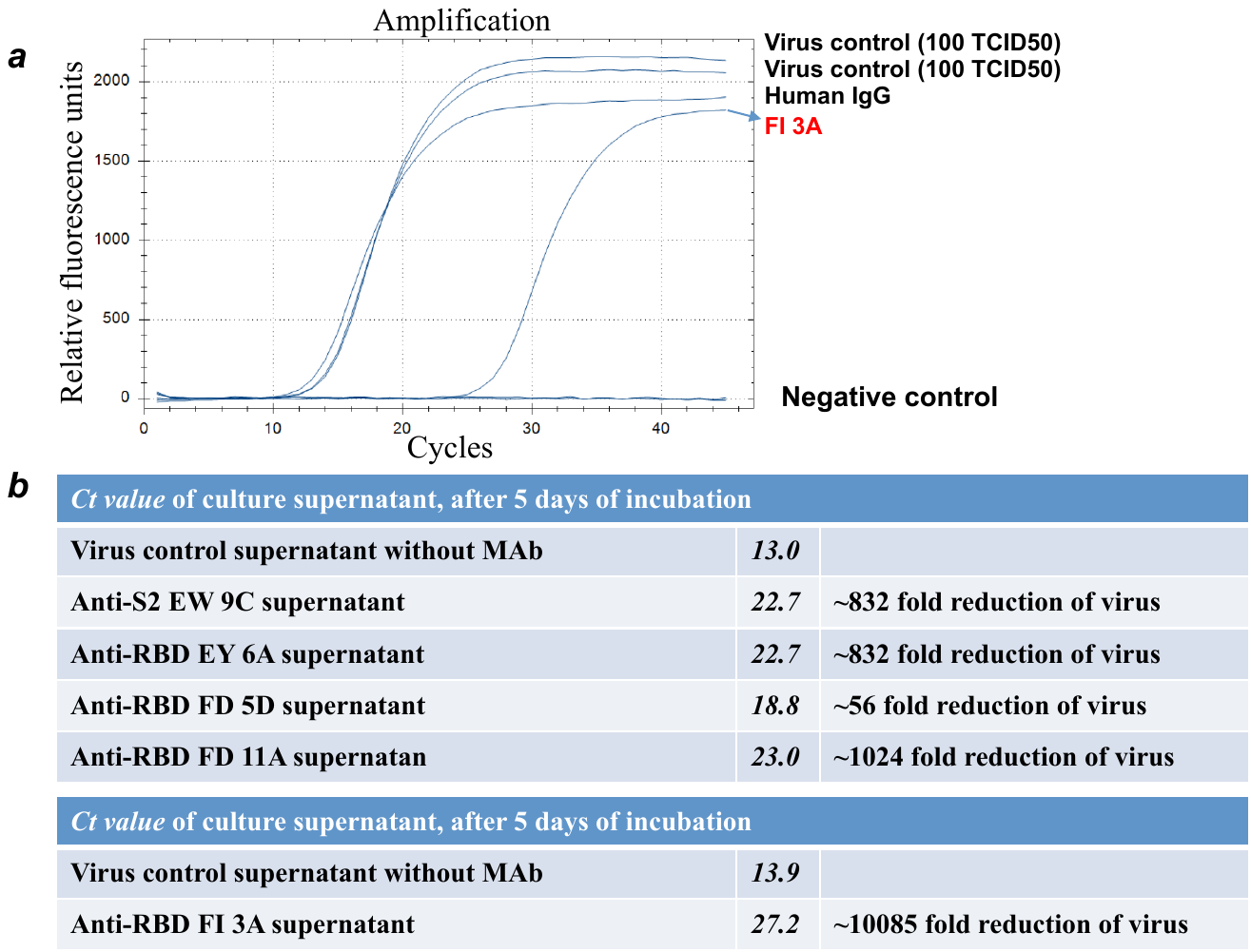

**Supplemental Figure 5. Neutralisation of SARS-CoV-2 by monoclonal antibodies in the PCR-based neutralisation assay. (a)** An illustrative example of measuring Ct value of virus signal in the tissue-culture supernatant of SARS-CoV-2 infected Vero E6 cells using an E gene-based real-time reverse-transcription PCR assay. The right shift of the amplification plot reflects the increase in Ct value and the decrease of viral load. **(b)** Neutralisation data for MAbs EW 9C, EY 6A, FD 5D, FD 11A and FI 3A. Increases in Ct value indicate decreases in virus loads. Each unit increase indicates a 2x reduction resulting from the presence of MAb. A 10x increase in Ct = 1,024 fold reduction of virus load.

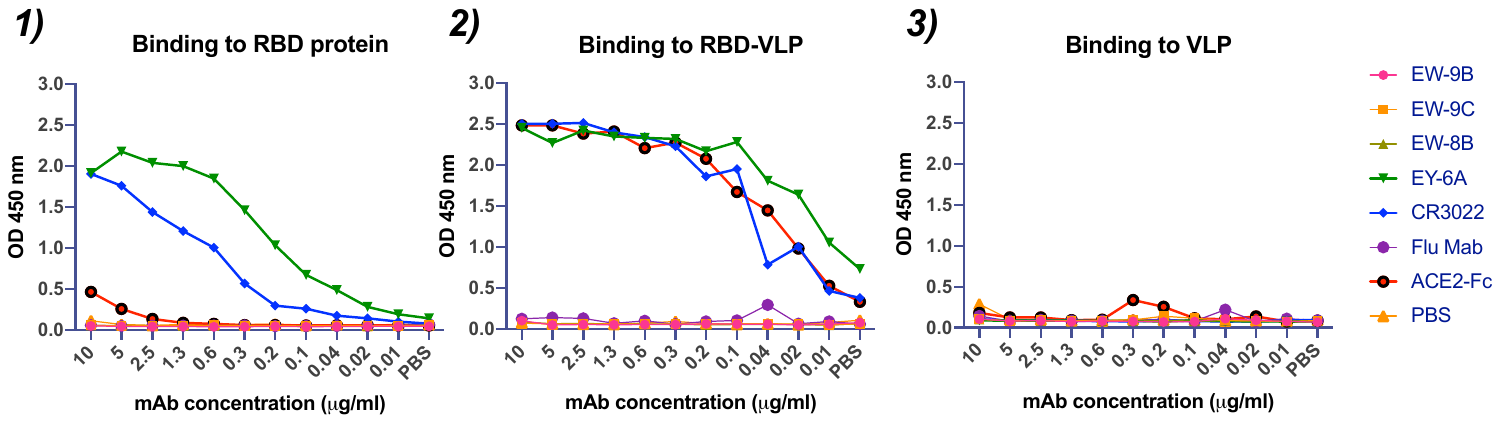

**Supplemental Figure 6. Binding of ACE2-Fc and MAbs to ST1-RBD (340-538) compared to ST1-RBD-mi3VLP in the ELISA.** RBD bound directly to plate (Panel 1) fails to bind ACE2-Fc, but RBD-VLP bound to plate (Panel 2) exposes the ACE2 binding site on RBD, and the epitopes bound by the MAbs CR3022 and EY 6A. VLP only (Panel 3) was included as a control in the assay. Other MAbs EW 9B, EW 9C, EW 8B bind elsewhere on the spike glycoprotein (therefore are negative in this assay), anti-influenza H7 haemagglutinin MAb is a negative control. Each experiment was repeated twice.

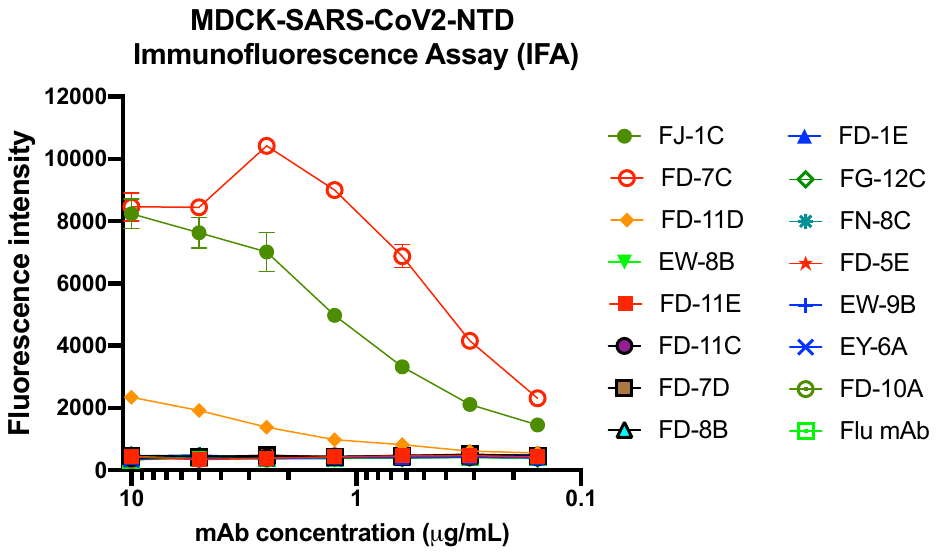

**Supplemental Figure 7. The binding activity of anti-SARS-CoV-2 NTD with MDCK-NTD cells in immunofluorescence.** Anti-influenza neuraminidase Z3-B2 (Flu MAb) was included as control in the experiment. Each experiment was repeated twice. Values are presented as mean ± standard error of the mean.

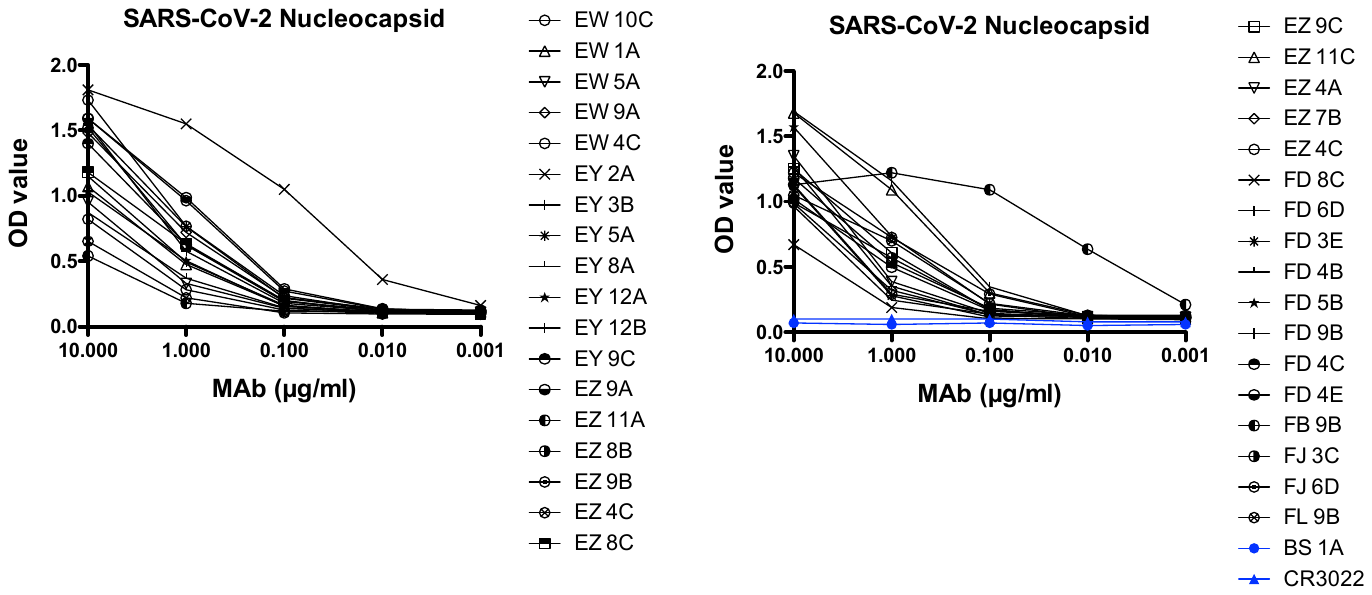

**Supplemental Figure 8. The binding activity of anti-SARS-CoV-2 nucleocapsid (N) antibodies with antigens of SARS-CoV-2 in ELISA.** Anti-influenza H3 BS-1A and anti-SARS spike CR3022 MAbs were included as controls. Each experiment was repeated twice. OD450 values are presented as mean ± standard error of the mean.

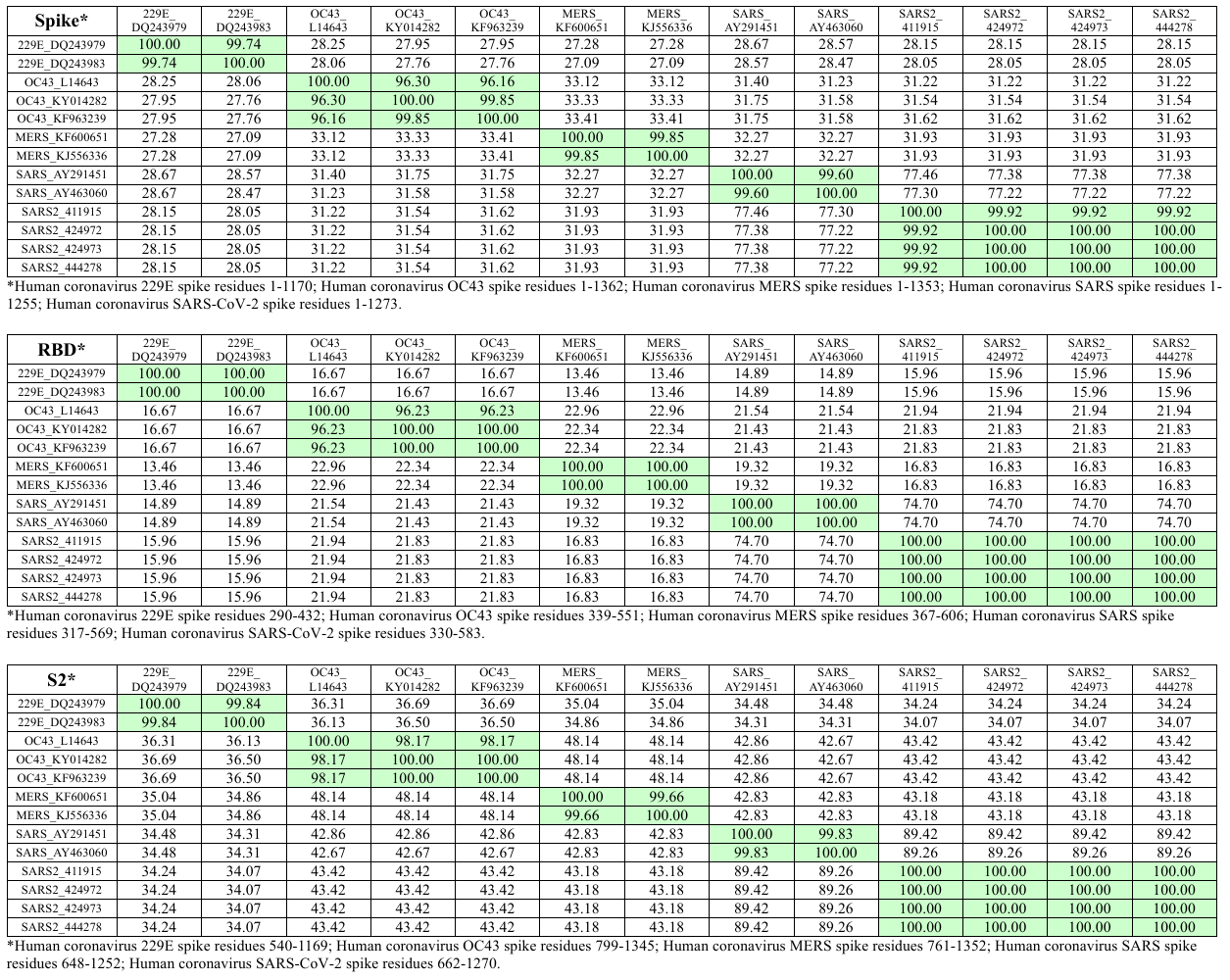

**Supplemental Figure 9. Percentage identities of spike glycoprotein, RBD and S2 amino acid residues among human coronaviruses.** Sequences were retrieved from the Genbank database (DQ243979, DQ243983, KY014282, KF963239, L14643, KF600651, KJ556336, AY291451, AY463060) and the EpiFlu^TM^ database of GISAID (EPI_ISL_411915, EPI_ISL_424972, EPI_ISL_424973, EPI_ISL_444278).

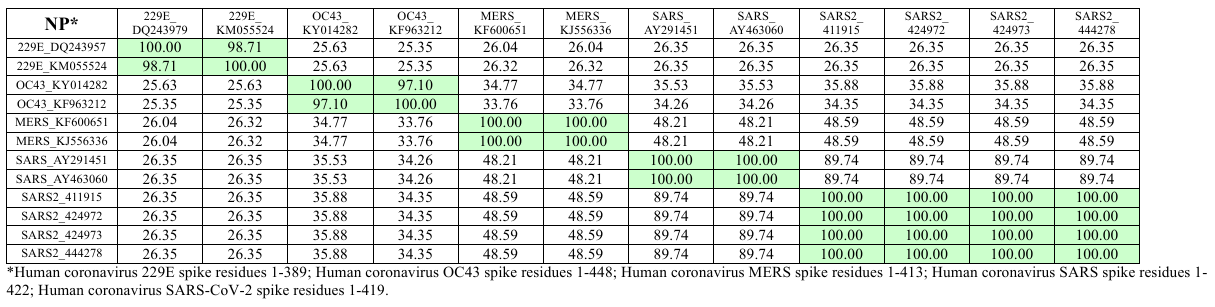

**Supplemental Figure 10. Percentage identities of nucleocapsid amino acid residues among human coronaviruses.** Sequences were retrieved from the Genbank database (DQ243957, KM055524, KY014282, KF963212, KF600651, KJ556336, AY291451, AY463060) and the EpiFlu^TM^ database of GISAID (EPI_ISL_411915, EPI_ISL_424972, EPI_ISL_424973, EPI_ISL_444278).
